# Aquaporins enriched in endothelial vacuole membrane regulate the diameters of microvasculature in hyperglycemia

**DOI:** 10.1101/2023.01.23.525218

**Authors:** Changsheng Chen, Yinyin Qin, Yidan Xu, Xiaoning Wang, Wei Lei, Xiaozhong Shen, Lixun Chen, Linnong Wang, Jie Gong, Yongmin Wang, Shijun Hu, Dong Liu

## Abstract

**Background:** In patients with diabetic microvascular complications, decreased perfusion or vascular occlusion, caused by reduced vascular diameter, is a common characteristic that will lead to insufficient blood supply. However, the regulatory mechanism remains undetermined and the treatment approach is lacking.

**Methods:** Gene expression of *Aquaporin 1* (*AQP1*) was compared between diabetic and non-diabetic human retina samples. Confocal live imaging analysis of zebrafish was utilized to investigate the hyperglycemia-induced alterations in vascular morphology. Transcriptomic and single-cell RNA sequencing, *in situ* hybridization, and quantitative PCR were used to analyze gene expressions. Gene knock-out zebrafish and stable transgenic lines were established for functional analysis. Human embryonic stem cells (H1 line), H1-derived endothelial cells (ECs), human umbilical vein endothelial cells (HUVECs), and human retinal microvascular endothelial cells (HRMECs) were used for functional analysis.

**Results:** Firstly, we found that the expression of human *AQP1* was downregulated in diabetic human retina samples (49 healthy vs. 54 diabetic samples) and high-glucose-treated human retinal microvascular endothelial cells. Then, we observed the reduction of vascular diameter and compromised perfusion in high glucose-treated zebrafish embryos. Two aquaporins (*aqp1a.1* and *aqp8a.1*), highly enriched in ECs, were significantly down-regulated under hyperglycemic conditions. *Aqp1a.1* and/or *aqp8a.1* loss-of-function leaded to reduction in intersegmental vessel diameters, recapitulating the phenotype of the hyperglycemic zebrafish model.

Overexpressing *aqp1a.1/aqp8a.1* in zebrafish ECs promoted the enlargement of microvascular diameters. Moreover, the reduced vessel diameters induced by high-glucose treatment was rescued by aquaporin overexpression. In addition, both *aqp1a.1* and *apq8a.1* were localized in the intracellular vacuoles in cultured ECs as well as the ECs of sprouting ISVs, and loss of Aqps caused the reduction of those vacuoles, which was required for lumenization. Loss of AQP1 had no influence on EC differentiation from human stem cells but strongly inhibited the vascular tube formation of differentiated ECs.

**Conclusions:** EC-enriched aquaporins regulate intracellular vacuole-mediated blood vessel diameter hyperglycemia. All of these suggest that EC-expressed aquaporins might be potential targets for gene therapy to cure diabetes-related vascular perfusion defects.

## Introduction

Hyperglycemia-caused microvascular complications, including hemorrhages, microaneurysms, capillary non-perfusion, and neovascularization ^1–5^, are the major source of morbidity and mortality in diabetes. Although the main therapeutic strategy in diabetes relies on glycemic control, targeting underlying factors contributing to microvascular complications is also necessary. During diabetic hyperglycemia, decreased vessel diameter is a common characteristic that will cause insufficient blood supply and aggravate vascular dysfunction ^6^. High blood glucose level is also able to cause fatty or other deposits to form inside blood vessels which narrow the vascular lumen and lessens blood flow. In development, the vascular lumen can be formed through vacuole coalescence theory, in which the intracellular vacuoles of endothelial cells (ECs) coalesce and gives rise to a seamless lumen ^7, 8^. However, it remains unclear which regulators are involved in regulating the blood vessel diameters under hyperglycemic conditions.

Aquaporin (AQPs) are a superfamily of tetrameric membrane proteins that involve in the transportation of water across cell plasma membranes ^9, 10^. Several of these small, integral membrane proteins are also known for the transfer of other small, uncharged solutes (e.g., glycerol) to maintain fluid homeostasis ^11^. Accumulating evidence suggests that specific classes of AQPs are involved in angiogenesis and EC function under physiological and pathological conditions ^12–14^. For instance, AQP1 is mainly expressed in microvascular endothelium and tumor microvessels, and it plays a role in driving water transport in lamellipodia and facilitating lamellipodial extension and cell migration ^13, 14^. Meanwhile, knocking-down AQP1 strongly inhibits the growth of new blood vessels, providing evidence that AQP1 is involved in both physiological and pathological angiogenesis ^15, 16^. Nevertheless, whether AQPs have a role in regulating vascular diameter is uncharacterized.

The identification of novel regulators involved in microvasculature diameter regulation in hyperglycemia will expand our knowledge of therapeutic strategy for diabetes-related microvascular complications. However, such study is generally hindered by the lacking of superior models for imaging the blood vessels at high resolution *in vivo*. To address this, the zebrafish (*Danio Rerio*) model was utilized in our study, which offers advantageous properties in cardiovascular research ^17, 18^. The transparency of zebrafish embryos allows us to perform high-resolution imaging analysis of vascular development *in vivo*. Importantly, the zebrafish can live up to the first week of development without a functional vasculature, allowing a detailed analysis even in animals with severe cardiovascular defects; by contrast, avian and mammalian embryos die rapidly in the absence of a functional cardiovascular system. Moreover, establishing zebrafish diabetes mellitus (DM) models are convenient and easy to manipulate ^19^.

In the current work, we found that human *AQP1* gene was down-regulated in diabetic retina samples and high-glucose treated human retinal microvascular endothelial cells. Under hyperglycemic state, zebrafish embryos exhibited narrowed and reduced perfusion of blood vessels in both eyes and trunks. Meanwhile, two endothelial-enriched aquaporins, *aqp1a.1* (orthologous to human *AQP1*), and *aqp8a.1*, were significantly down-regulated upon high glucose treatment. Further, we found that *apq1a.1* or *aqp8a.1* loss of function leaded to the reduction in vessel diameter and perfusion of intersegmental vessels, recapitulating the vascular phenotype of hyperglycemic zebrafish embryos. However, the gain-of-function of *apq1a.1* or *aqp8a.1* in zebrafish embryos caused the increased diameter of blood vessels. Moreover, overexpression of *apq1a.1* in hyperglycemic zebrafish embryos rescued the defective vascular phenotype. Both *in vivo* and *in vitro* experiments implied that zebrafish Aqps were able to regulate the size and number of intracellular vacuoles of ECs. All taken together, AQPs are important regulators in regulating blood vessel diameter under hyperglycemic conditions, and this specific role of AQPs might be exploited for clinical benefits by modulation of their expressions.

## Materials and Methods

### Study approval

Experiments involving human samples were approved by the Affiliated Hospital of Nantong University (2020-K013). Retina samples from 103 donors were obtained from Nanjing Red Cross Eye Bank (Nanjing, China) between 2019 and 2021, and informed written consent was obtained from each donor. The 49 non-diabetic retina donors included 31 males and 18 females (average age, 62; age range, 23-85 years), and the 54 diabetic donors included 39 males and 15 females (average age, 66; age range, 37-88 years). All animal experimentation was carried out in accordance with NIH Guidelines for the care and use of laboratory animals (http://oacu.od.nih.gov/regs/index.htm) and ethically approved by the Administration Committee of Experimental Animals, Jiangsu Province, China (Approval ID: SYXK(SU) 2017–111). Best efforts were made to minimize the number of animals used and prevent their suffering.

### Zebrafish husbandry and strains

The wild-type AB line and transgenic lines *Tg(kdrl:EGFP)* ^20^, *Tg(fli1ep:EGFP-CAAX)^ntu^*^666^, *Tg(fli1ep:aqp1a.1-mCherry)^ntu^*^667^, *Tg(fli1ep:aqp8a.1-EGFP)^ntu^*^668^, and *Tg(fli1:EGFP– cdc42wt)* ^7^ were used in this study. All zebrafish embryos and adult fishes were raised and maintained at 28.5 °C.

### Construction of transgenic lines

*Tg(fli1ep:EGFP-CAAX)^ntu^*^666^ transgenic line was generated by injecting *pTol2-fli1ep-EGFP-CAAX* into the progeny of wild-type AB line together with Tol2 transposase mRNA into 1-cell stage wild-type fertilized eggs (1 ng/embryo) according to the Tol2kit protocol ^21^. *Tg(fli1ep:aqp1a.1-mCherry) ^ntu^*^667^ and *Tg(fli1ep:aqp8a.1-mCherry) ^ntu^*^668^ transgenic lines were generated by using the same strategy (Fig.S1).

### High glucose treatment

The zebrafish embryos were incubated in high glucose solution (4% D-glucose w/v) for 24 hours from 48 hpf and imaged at 72 hpf. The embryos incubated in the E3 medium were used as control. To measure the glucose level in zebrafish embryos after glucose treatment, the embryos were first sacrificed with rapid chilling (submerged fish embryos into 2-4 °C chilled water). Following confirmation of the absence of heartbeat under a stereo microscope, the embryos were rinsed with E3 medium twice and then dried with a paper towel and placed into 2-ml microcentrifuge tube. One-hundred embryos were collected for each time point (48, 60, 72, 84, and 96 hpf, respectively) and divided into 5 groups (5 biological replicates). The embryos were homogenized and centrifuged to collect the supernatant. Two microliters of the supernatant were placed on a glucometer strip (One-Touch Ultra 2), and the total free glucose level was measured with a glucometer (One-Touch Ultra 2).

### ECs sorting, RNA extraction, RNA-seq, and qPCR

To isolate ECs, the control and high-glucose treated *Tg(kdrl:EGFP)* embryos (300-400 embryos for each) were collected at 96 hpf, and followed by washing with PBC three times and digested with 0.25% trypsin at 37°C. The digested cells were collected by centrifugation and allowed to pass through a 40 mm FACS tube (BD Falcon, 352340). The EGFP-positive cells were sorted by fluorescence-activated cell sorting on FACS Aria3 (BD Biosciences). Total mRNAs of the retina samples, zebrafish embryos, or ECs were extracted with TRIzol Reagent (Invitrogen, USA) and quantified and qualified by NanoDrop (ThermoFisher Scientific, USA) and 1% agarose gel. To study gene expression changes after high glucose treatment, RNA-seq was performed with the prepared mRNAs from 72 and 96 hpf-control and high glucose-treated zebrafish embryos. Three independent replicates of the samples were analyzed for each treatment. All RNA samples were submitted to GENEWIZ Science (Suzhou, China), and deep sequencings were performed on an Illumina Hiseq2500. Quantitative RT-PCR was conducted in a total 20 µl reaction volume with 10 µl SYBR premix (TIANGEN). The relative RNA amounts were calculated with the comparative CT (2-DDCT) method and normalized with *ef1a* and *GAPDH* as the references for zebrafish and human retina samples, respectively. The primers for qPCR are listed in Supplementary Table S1.

### Single-cell preparation and gene expression profile analysis

Zebrafish embryos (*Tg(fli1ep:EGFP-CAAX)^ntu^*^666^) were collected and dissociated with 2.5% trypsin. Single cells were captured and processed for RNA-seq using the Chromium platform (10x Genomics). Cell Ranger 3.0.2 (https://github.com/10XGenomics/cellranger) was utilized to process and de-multiplex the raw sequencing data to a single-cell level gene counts matrix. An analysis of the single-cell RNA-seq data was done with the package Seurat 3.0 (https://satijalab.org/seurat/install.html) ^22^. A Seurat object was created by filtering cells with the number of unique expressed genes in a range of 500-3,000, and less than 5% of counts were mapping to the mitochondrial genome and the filtering genes that were expressed in at least 3 single cells. Additionally, an R package sctransform was applied to remove technical variation ^23^. Dimensionality reduction was performed on the entire dataset based on t-distributed stochastic neighbor embedding (t-SNE) analysis, and visualization of specific gene expression patterns across groups on t-SNE and violin plots were performed using functions within the Seurat package.

### Whole-mount in situ hybridization (WISH)

WISH with antisense RNA probes was performed as described previously ^24^. Templates for making probes to detect the expression of *aqp1a.1* (NM_207059.1) and *aqp8a.1* (NM_001004661.1) was cloned from cDNA fragments. Primers for WISH were listed in Supplementary Table S1. After hybridization, images of the embryos were acquired with an Olympus stereomicroscope MVX10 equipped with an Olympus DP71 camera.

### Generation of *aqp1a.1* or *aqp8a.1* knock-out mutants

To generate *aqp1a.1* or *aqp8a.1* gene mutant zebrafish, we used a CRISPR/Cas9-mediated approach. CRISPR/Cas9 target sites were designed to identify the sequences in the second exon of *aqp1a.1* or *aqp8a.1*. Primers for single-RNA (sgRNA) synthesis were listed in Supplementary Table S1. Each sgDNA was individually subcloned into a pT7 vector and transcribed into sgRNA in vitro using the T7 mMessage mMachine kit (Ambion). Microinjection was performed with 1-cell stage zebrafish embryos, and each embryo was co-injected with 100 pg sgRNA and 200 pg Cas9 mRNA. G0 generations were examined by PCR and subsequently Sanger sequencing. A mutant with 7 bp deletion at the second exon of *aqp1a.1,* which leads to the occurrence of a stop codon downstream of the Cas9 cutting site, was identified. The heterozygous *aqp1a.1^+/-^* mutants were incrossed to obtain the homozygous F2 progenies.

### Morpholino-mediated gene knock-down of *aqp1a.1* or *aqp8a.1* in zebrafish

The gene-specific morpholinos (Gene Tools, LLC) were used to block the splicing of the pre-mRNA. Morpholino antisense oligomers (referred to as MO) were prepared according to the manufacturer’s instructions. The MO sequences used are listed in Supplementary Table S1. In this experiment, 5 ng morpholino oligo was microinjected into the 1-cell stage zebrafish embryo.

### HgCl_2_ treatment

Functional inhibition of Aqp1a.1 and Aqp8a.1 using HgCl_2_ was performed with dechorionated embryos at 22 hpf in the E3 medium. Three different HgCl_2_ concentrations (0.2, 0.4, and 1.6 μM) were used. The embryos were incubated in the HgCl_2_-containing E3 medium from 22 hpf and imaged at 48 hpf.

### Microangiography and confocal imaging

For confocal imaging of blood vessel development, zebrafish embryos were treated with 1-phenyl-2-thiourea (Sigma Aldrich) to inhibit the pigmentation. After manually dechorionation, embryos were anesthetized with egg water/0.16 mg/ml tricaine/1% 1-phenyl-2-thiourea (Sigma) and embedded in 0.6% low melting agarose. For microangiography, 5 μg/μl Dextran TexasRed (Invitrogen) was injected into the common cardinal vein (CCV) at 72 hpf and immediately processed for optical imaging. Living imaging was performed with Nikon A1R confocal microscopy.

### Quantitative analysis of blood vessel diameter, vascular perfusion, and intracellular vacuoles

The vessel diameters were quantified in Fiji using the Vessel Analysis plugin ^25^. For zebrafish, 5 locations on IOC, DA, and PCV were selected and measured, and each measurement was used as a data point. For zebrafish ISV diameter measurement, 5 ISVs beyond the yolk yac extension were selected, and an individual ISV was measured at 5 positions. Five fish were applied in each group for analysis. The average of these data points was statistically used as the diameter of the analyzed vessel. For human retinal vessels, measurements were carried out with fluorescein angiograms. The diameters of the central retinal vessels in the optic disk and the branching retinal vessels were respectively measured and quantified. The vascular perfusion analysis in zebrafish was based on the microangiographic images. Completely or partially lost the red fluorescence in ISV was defined as defective-perfused ISV. The area of intracellular vacuoles in zebrafish ISV was quantified with the transgenic line *Tg(fli1:EGFP–cdc42wt)* by using Fiji. Five fishes were applied in each group, and 5 ISVs beyond the yolk yac extension were selected for analysis.

### Cell culture and transfection

Cell culture and maintenance of human ESCs (H1 line) were performed as previously described ^26^. The cells were cultured on Matrigel-coated plates (ESC qualified, BD Biosciences, San Diego, CA) using hESC mTeSR-1 cell culture medium (StemCell Technologies, Vancouver, Canada) under conditions of 37 °C, 95% air, and 5% CO2 in a humidified incubator. Human ECs isolated from 2 donors were conducted as described previously ^27^. Human umbilical vein endothelial cells (HUVECs, Cell System) were cultured in endothelial cell growth medium according to the protocol provided by the manufacturer (VascuLife, Cell System). Lipofectamine™ 3000 Transfection Reagent (ThermoFisher Scientific) was used for plasmid transfection. The *aqp1a.1* or *aqp8a.1* cDNA was subcloned into pmCherry-N1 or pEGFP-N1 vector (Addgene). Human retinal microvascular endothelial cells (HRMECs, Cell System) were cultured in a complete classic medium supplemented with/without high-content glucose (50 or 100 mM). Cells were incubated in normal medium and high glucose medium for 48 h and subjected to RNA extraction.

### Generation of AQP1-knockout H1 cell lines with epiCRISPR system

Genome editing with the epiCRISPR system was performed as previously described ^26^. 4×10^5^ human ESCs (H1 line) were disassociated and cultured in a 6-well plate with mTeSR-1 cell culture medium for 24 hours. On day 1, 2 µg of the epiCRISPR plasmid were transfected into cells by using lipofectamine 3000 (Life Technologies). On day 2, cells were selected by puromycin (0.2–0.5µg/ml). On day 5, day 10, and day 15, genomic DNA was extracted from cells and the gRNA targeting sites were amplified using PCR. The PCR products were purified and digested with restriction enzymes. For analysis of single cell-derived clones, the cells were disassociated into single cells at 15 days post-transfection, and seeded them onto the Matrigel-coated plates with puromycin-free mTeSR-1 medium for 15 days. Individual colonies were picked and genotyped.

### EC differentiation

Embryoid body (EB)-based EC differentiation method was performed as described previously ^28^. For EC differentiation, EBs were formed in an ESC medium depleted of recombinant human FGF-2 (rhFGF-2) (R&D Systems) on day 0. On day 1, the medium was supplemented with 20 ng/ml Activin A (R&D Systems) and 20 ng/ml BMP4 (R&D Systems). On day 2, the medium was supplemented with 10 ng/ml rhFGF-2, 20 ng/ml Activin A, and 20 ng/ml BMP4. On day 4, the EBs were seeded onto Matrigel-coated dishes, and the medium was supplemented with 10 ng/ml rhFGF-2 and 20 ng/ml VEGF (R&D Systems) for EC expansion. The EBs were harvested by Accutase (Sigma-Aldrich) and sorted using an anti-CD31 monoclonal antibody (BD Biosciences, catalog 558068) on day 0, day 1, day 2, day 5, and day 10, respectively. The differentiation efficiency was calculated by the number of CD31+CD144+ cells divided by the number of total differentiated cells, excluding dead cells at the end of induced differentiation.

### Statistical analysis

Statistical analysis employed a two-tailed, unpaired Student’s t-test, One-way ANOVA, or Fisher’s exact test is described in figure legends. The chi-square test (χ2-value) was used for the intergroup comparison of categorical data. All data are presented as mean ± s.e.m. P<0.05 was considered to be statistically significant.

## Results

### Expression of *AQP1* in human retina specimens and cultured HRMECs

A total of 103 donors (n=54 in the diabetic group and n=49 in the non-diabetic group) were enrolled. Fig.1A showed the representative fundus images and fluorescein angiograms of non-diabetic retinas and diabetic subjects with different grades of DR, starting from mild non-proliferative diabetic retinopathy (NPDR) to severe proliferative diabetic retinopathy (PDR). The mean diameters of central retinal vessels (central retinal artery and vein) and the retinal vessel branches (retinal arterioles and venules) were measured (Fig.1B). There was no difference in the central retinal vessel between diabetic and non-diabetic retina samples, whereas the diameter of the retinal vessel branches was significantly decreased in diabetic retinas (Fig.1C and D). Next, the mRNAs of both groups were isolated and applied to quantitative PCR. The statistical results demonstrated that the relative expression of *AQP1* was significantly lower in diabetic samples than in non-diabetic samples (Fig.1E). The percentages of donors with downregulated *AQP1* were 77.8% (42/54) and 42.9% (21/49) in diabetic and non-diabetic groups, respectively (Table 1). The *AQP1* expression was negatively correlated with diabetes (P=0.0003). Moreover, there was no difference between the groups in terms of age and sex (P=0.3983 and P=0.3439, respectively) (Table 2). In the diabetic group, 19 patients were diagnosed with NPDR, and 35 patients were diagnosed with PDR. However, there was no *AQP1* expression difference between NPDR patients (16 samples showed down-regulated *AQP1* expression) and PDR patients (26 samples showed down-regulated *AQP1* expression). The expression of *AQP1* in endothelial cells under high glucose conditions was further assessed with cultured human retinal microvascular endothelial cells (HRMECs) (Fig.1F). Compared with cells cultured in a complete classic medium, high glucose-treated cells displayed a lower *AQP1* expression (Fig.1G). These results suggested that the endothelial AQP1 was negatively correlated with diabetes or hyperglycemia.

**Figure 1.**
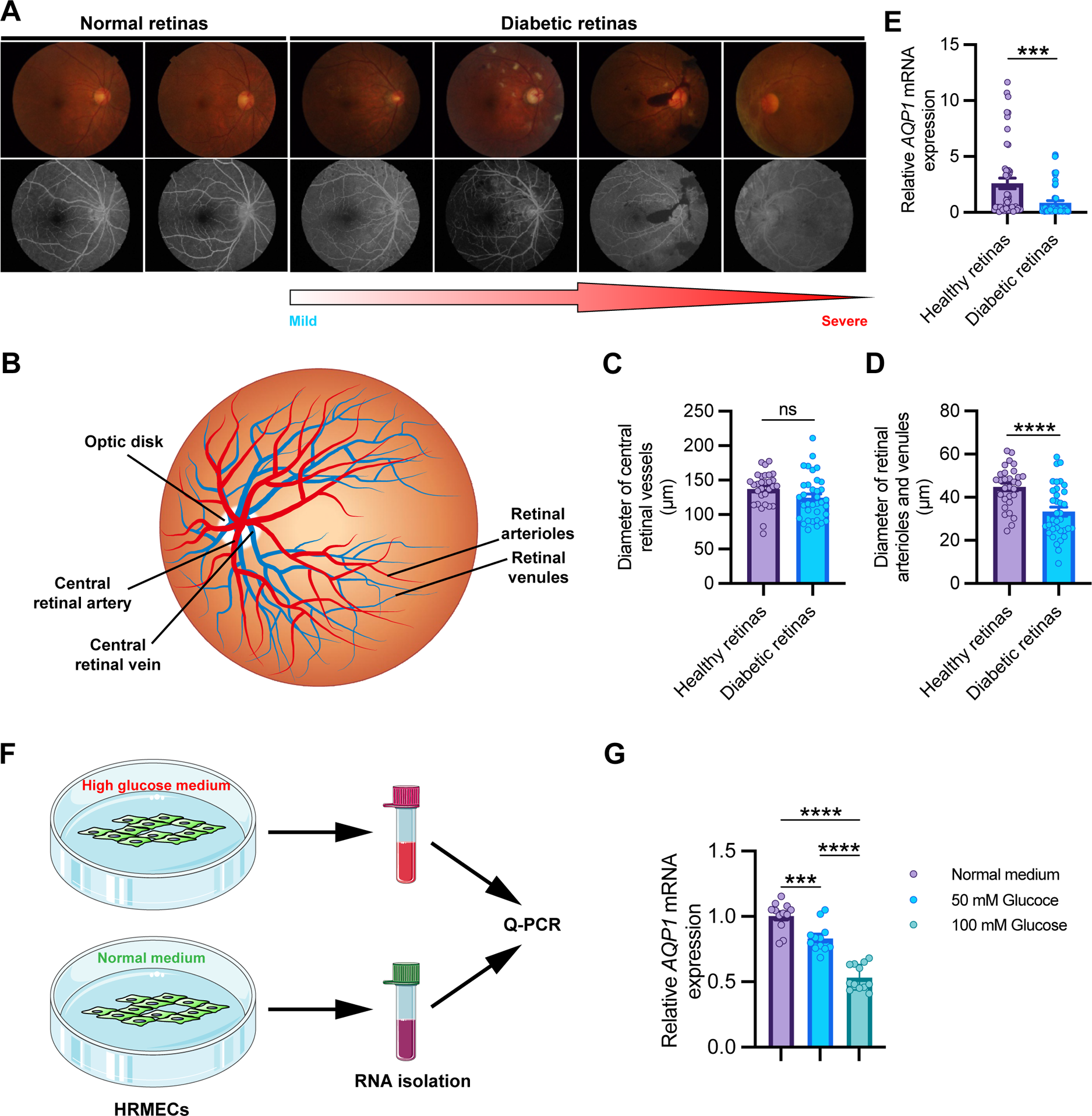
*AQP1* expression is downregulated in retina samples from diabetic patients. **A,** Representative fundus images (upper) and fluorescein angiography (lower) of normal retinas and retinas with diabetic retinopathy (DR) at different stages (from mild to severe). **B,** Representative sketch of human retinal vessels. **C and D,** Diameters of the central retinal vessels (central retinal artery and vein) and the branching retinal vessels (retinal arterioles and venules). Each data point represents the mean diameter of central retinal vessels in C, and the mean diameter of the branching retinal vessels in D. Data are shown as mean ± s.e.m. A two-tailed, unpaired Student’s t-test is applied. ns, not significant. ****, p < 0.0001. **E,** Quantification of *AQP1* expression in healthy (*n=49*) and diabetic (*n=54*) retina samples. Each data point represents the *AQP1* expression in an individual sample. Data are shown as mean ± s.e.m. A two-tailed, unpaired Student’s t-test is applied. ***, p < 0.001. **F,** Schematic representation of cell experiment. HRMECs were cultured in normal and high glucose mediums, respectively. The RNAs of each group were separately isolated and quantified for *AQP1* expression. **G,** Quantification of *AQP1* expression in HRMECs cultured in normal and high glucose medium. One-way ANOVA analysis is applied. ****, p < 0.0001.

**Table 1.** Differential expression of *AQP1* mRNA in non-diabetic and diabetic retinas

| Type | number | Relative expression |  | Negative | P |
| --- | --- | --- | --- | --- | --- |
|  |  | <0.5<sup>a</sup> | >0.5 |  |  |
| Retinas from non-diabetic donors<br>(Healthy subjects) | 49 | 21 | 28 | 42.9% | 0.0003 |
| Retinas from diabetic donors<br>(Diabetic subjects) | 54 | 42 | 12 | 77.8% | $\chi^2=13.190$ |

<sup>a</sup>Two-folds downregulated.

**Table 2.** Correlation of AQP1 expression with age and sex

| Group |  | number | Relative expression |  | Negative | P |
| --- | --- | --- | --- | --- | --- | --- |
|  |  |  | <0.5<sup>a</sup> | >0.5 |  |  |
| Age | ≤64<sup>b</sup> | 50 | 28 | 22 | 56.0% | 0.3983 |
| | >64 | 53 | 34 | 19 | 64.1% | $\chi^2=0.7143$ |
| Sex | Male | 70 | 45 | 25 | 64.3% | 0.3439 |
| | Female | 33 | 18 | 15 | 54.5% | $\chi^2=0.8958$ |

<sup>a</sup>Two-folds downregulated; <sup>b</sup>Mean age.

### Hyperglycemia leads to reduced vascular diameters in zebrafish model

To examine the effects of high glucose on microvasculature, a hyperglycemia zebrafish model was established using the transgenic line *Tg(fli1ep:EGFP-CAAX)^ntu^*^666^, in which an endothelial enhancer/promoter fragment (fli1ep) was employed to drive the specific expression of the enhanced green fluorescent protein (EGFP) in ECs, and the CAAX membrane targeting motif was able to guarantee the localization of EGFP at the surface of ECs (Fig.S1A). In this line, direct non-invasive visualization of the zebrafish vasculature can be achieved under conventional fluorescence microscopy or confocal microscopy. A hyperglycemic state in zebrafish embryos was achieved by immersing them into a high glucose solution (4% D-glucose w/v) from 2 days post fertilization (dpf), and fluorescence imaging of the vasculature was performed at 3 dpf (Fig.2A). The average total free glucose level in zebrafish embryos exposed to high glucose solution reached more than twice higher compared to PBS-treated controls (Fig.S2). The high glucose treatment did not yield phenotypical differences and developmental defects in zebrafish embryos (Fig.S3), but the vasculature was substantially influenced. Under this condition, the vessel diameters were significantly decreased compared with the untreated control counterparts, including the inner optic circle (IOC) in the eyes as well as the intersegmental vessels (ISVs), dorsal aorta (DA), and posterior cardinal vein (PCV) in the trunk (Fig.2B-C’’ and Fig.2E-F’’). The IOC in zebrafish eyes and the ISVs in zebrafish trunk were declined by 27.9% and 36.8%, respectively (Fig.2D and H). Also, microangiography analysis revealed that 59% of the narrowed ISVs were not perfused (Fig.2G), suggesting that the dysfunctional vascular lumens were formed after high glucose induction. In addition, exposing zebrafish embryos to 4% sucrose had no obvious influences on development and blood vessel diameters, excluding the potential effects posed by increased osmotic strength and viscosity in the media (Fig.S3).

**Figure 2.**
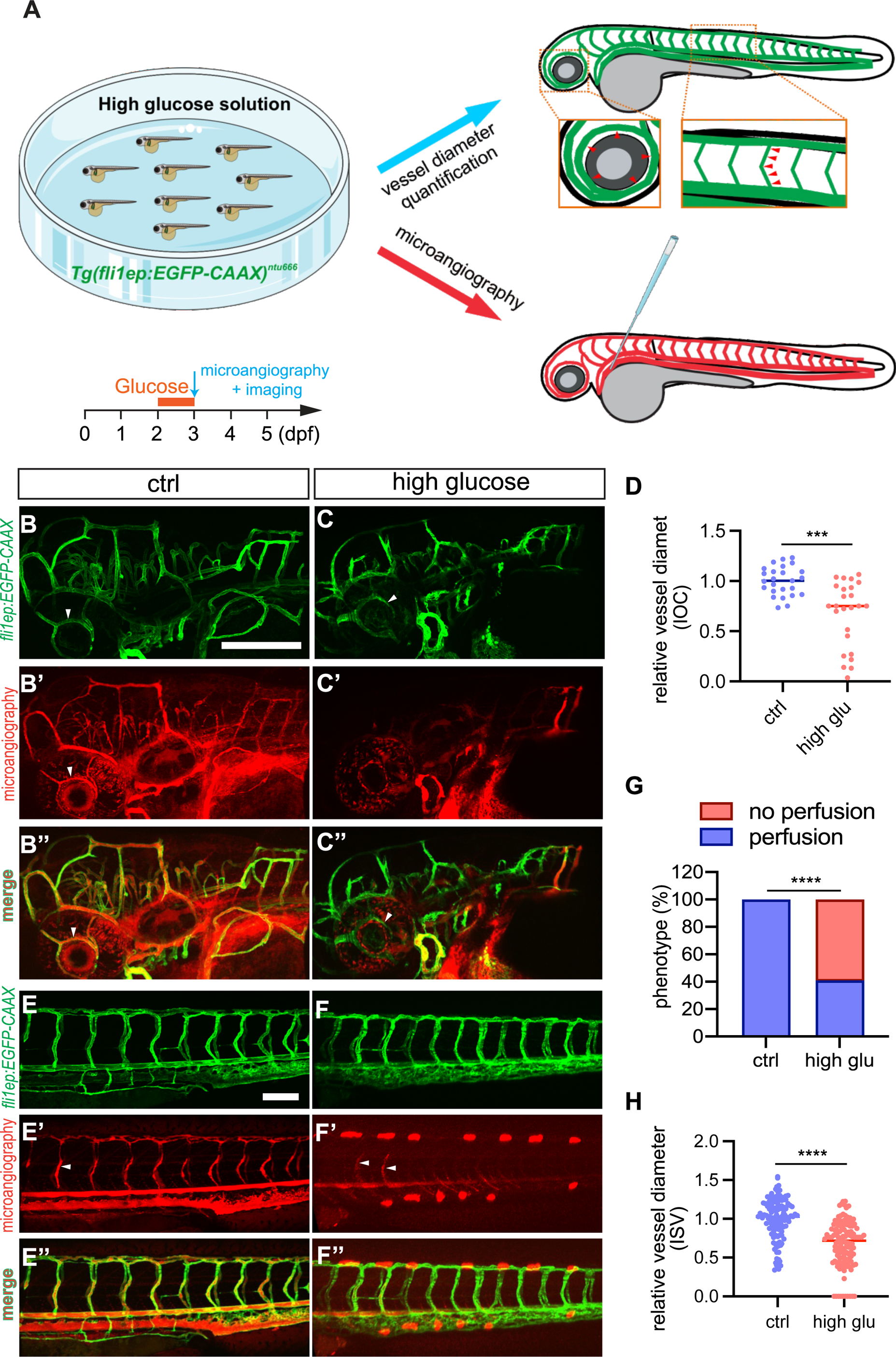
High glucose treatment affects vascular diameter in zebrafish. **A,** Schematic representation of establishing zebrafish hyperglycemia/DM model. Zebrafish embryos are immersed in high glucose solution (4% D-glucose w/v) from 2 to 3 dpf. Microangiography and confocal imaging are performed afterward. Red arrowheads indicate the positions for vessel diameter measurement on IOC or ISV. **B-C’’,** Representative confocal images of IOC phenotypes in 3-dpf *Tg(fli1ep:EGFP-CAAX)^ntu^*^666^ transgenic line with (*n=22*) or without (*n=14*) glucose treatment. White arrowheads indicate the IOC. **D,** Quantification of IOC diameter. **E-F’’,** Representative confocal images of ISV phenotypes in 3-dpf *Tg(fli1ep:EGFP-CAAX)^ntu^*^666^ transgenic line with (*n=22*) or without (*n=14*) glucose treatment. White arrowheads indicate the ISV. **G,** The incidence of normal and non-perfused ISVs in 3-dpf *Tg(fli1ep:EGFP-CAAX)^ntu^*^666^ transgenic line with (*n=22*) or without (*n=14*) glucose treatment. **H,** Quantification of ISV diameter. Each data point in **D** and **H** represents an individual vessel diameter measurement of IOC or ISV. Five fishes are analyzed for control and high glucose treatment. Data are shown as mean ± s.e.m. A two-tailed, unpaired Student’s t-test is applied. ***, p < 0.001. ****, p < 0.0001. Scale bars, 100 μm.

### High glucose treatment caused the downregulation of the expression of *aqp1a.1* and *aqp8a.1*

To further explore which genes are altered upon glucose treatment, whole-transcriptome sequencing, and single-cell transcriptome sequencing were performed. Compared to those in control zebrafish, the expression of a serial of genes were significantly altered at 72 and 96 hours post fertilization (hpf) embryos treated with high glucose, with 642 genes upregulated and 1,149 genes downregulated at 72 hpf, as well as 1,259 genes upregulated and 1,851 genes downregulated at 96 hpf (Fig.3A and B). Several EC-enriched genes, like *cxcr4a* and *esm1*, were found to be upregulated after high glucose treatment, while two aquaporins, *aqp1a.1,* and *aqp8a.1*, exhibited reduced expressions (Fig.3C and D). Further whole-mount ISH assay demonstrated that the zebrafish *aqp1a.1* and *aqp8a.1* were highly enriched in ECs of the vascular system (Fig.S4).

**Figure 3.**
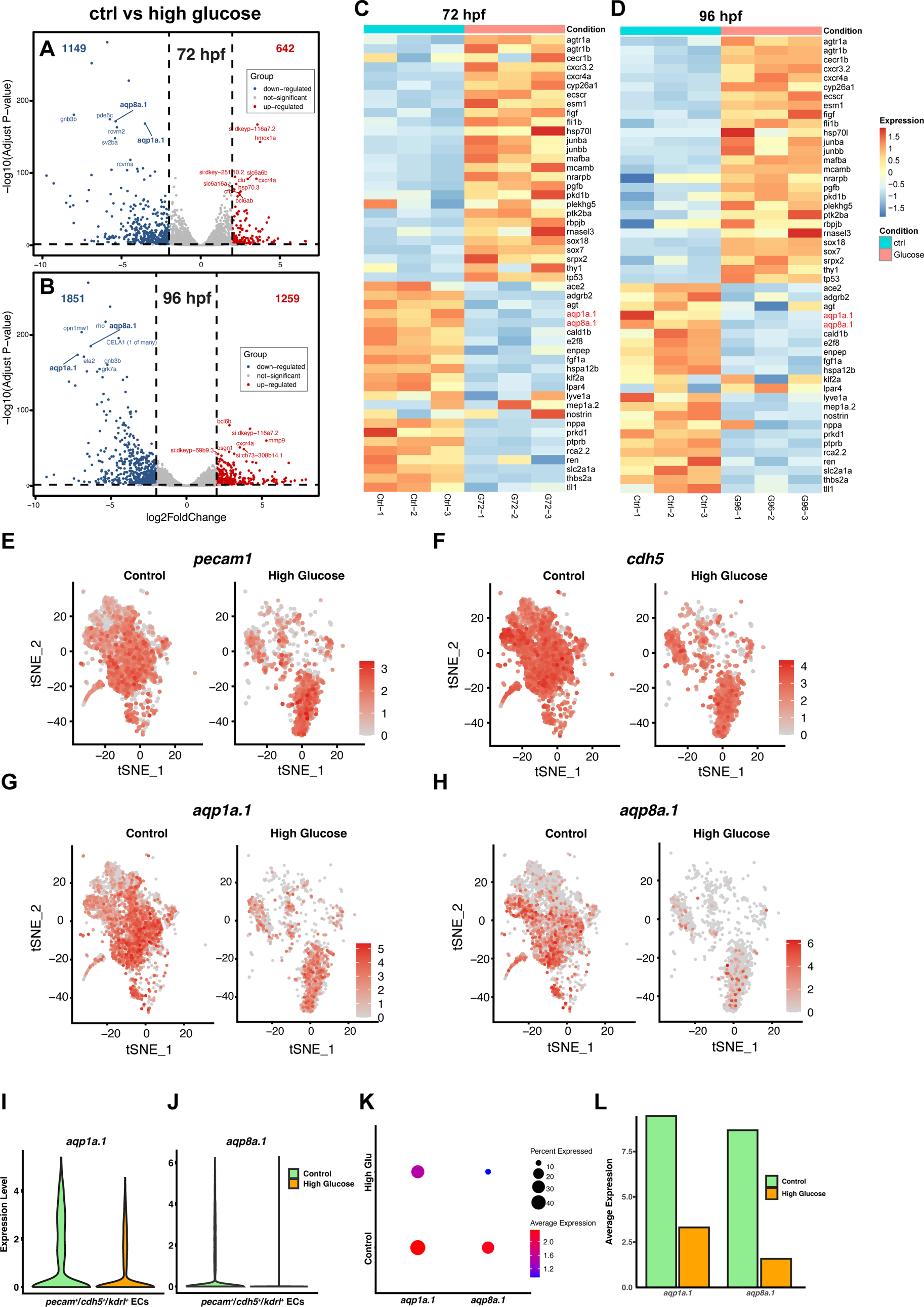
Whole-genome and single-cell transcriptomic profilings of *Tg(fli1ep:EGFP-CAAX)^ntu^*^666^ transgenic line with or without glucose treatment. **A and B**, Volcano plots of downregulated genes (blue dots), upregulated genes (red dots), and unchanged genes (grey dots) in the high glucose-treated group versus the control group at 72 and 96 hpf, respectively. **C and D,** Heatmaps based on bulk RNA-seq data represent the expression changes of the vascular-related DEGs. Zebrafish *aqp1a.1* and *aqp8a.1* are downregulated upon high glucose treatment. E-H, *t*-SNE plots demonstrate the relative distributions of *pecam1*, *cdh5*, *aqp1a.1*, and *aqp8a.1* across *pecam1^+^*/*cdh5^+^*/*kdrl^+^* cells in control and high glucose-treated groups. **I and J,** Expression levels of *aqp1a.1*, and *aqp8a.1* across *pecam1^+^*/*cdh5^+^*/*kdrl^+^* cells in control and high glucose-treated groups. **K and L,** Percentages and average expressions of *pecam1^+^*/*cdh5^+^*/*kdrl^+^* cells expressed *aqp1a.1*, and *aqp8a.1*, respectively.

Next, single-cell transcriptomic sequencing analysis was conducted to compare the transcriptional profiles of endothelial cells isolated from high glucose-treated and control embryos (*Tg(fli1ep:EGFP-CAAX)^ntu^*^666^). The EGFP-labeled ECs were dissociated and subjected to single-cell RNA-seq analysis using the Chromium (10x Genomics) platform (Fig.S5A). A total of 12,425 cells (8,217 and 4,208 cells for control and high glucose groups, respectively) were obtained and subjected to *t*-distributed stochastic neighbor embedding (*t*-SNE) projection (Fig.S5B). A cluster of cells that are positive for platelet EC adhesion molecule (*pecam-1*; *CD31*), cadherin 5 (*cdh5*; *CD144*), and kinase insert domain receptor-like (*kdrl*; *VEGFR2*) were separated and defined as ECs (Fig.S5C). A total of 6,896 *pecam1^+^*/*cdh5^+^*/*kdrl^+^* ECs (4,838 and 2,058 cells for control and high glucose groups, respectively) were subjected to *t*-distributed stochastic neighbor embedding (*t*-SNE) projection (Fig.3E-H, and Fig.S5C). High glucose treatment had no obvious effects on the expressions of *pecam-1* and *cdh5* (Fig.3E and F). Although *aqp1a.1* and *aqp8a.1* were also expressed across all separated ECs, their expressions were dramatically decreased under high glucose conditions (Fig.3G, H, I, and J). Meanwhile, the number of ECs that expressed *aqp1a.1*/*aqp8a.1* and the average expression of *aqp1a.1*/*aqp8a.1* in ECs sharply dropped after high glucose treatment (Fig.3K and L).

Additionally, the expressions of *aqp1a.1* and *aqp8a.1* in hyperglycemic zebrafishes were validated by real-time qPCR and ISH assay, and the results were consistent with the transcriptomic data (Fig.4A-H). To further confirm that *aqp1a.1* and *aqp8a.1* were down-regulated in ECs of high glucose-treated embryos, we performed the qPCR assay with isolated ECs. The ECs in the control and high glucose-treated group at 96 hpf were respectively sorted by FACs and subsequently used for RNA isolation (Fig.4I). Upon high glucose induction, both *aqp1a.1* and *aqp8a.1* in ECs were significantly downregulated (Fig.4J and K).

**Figure 4.**
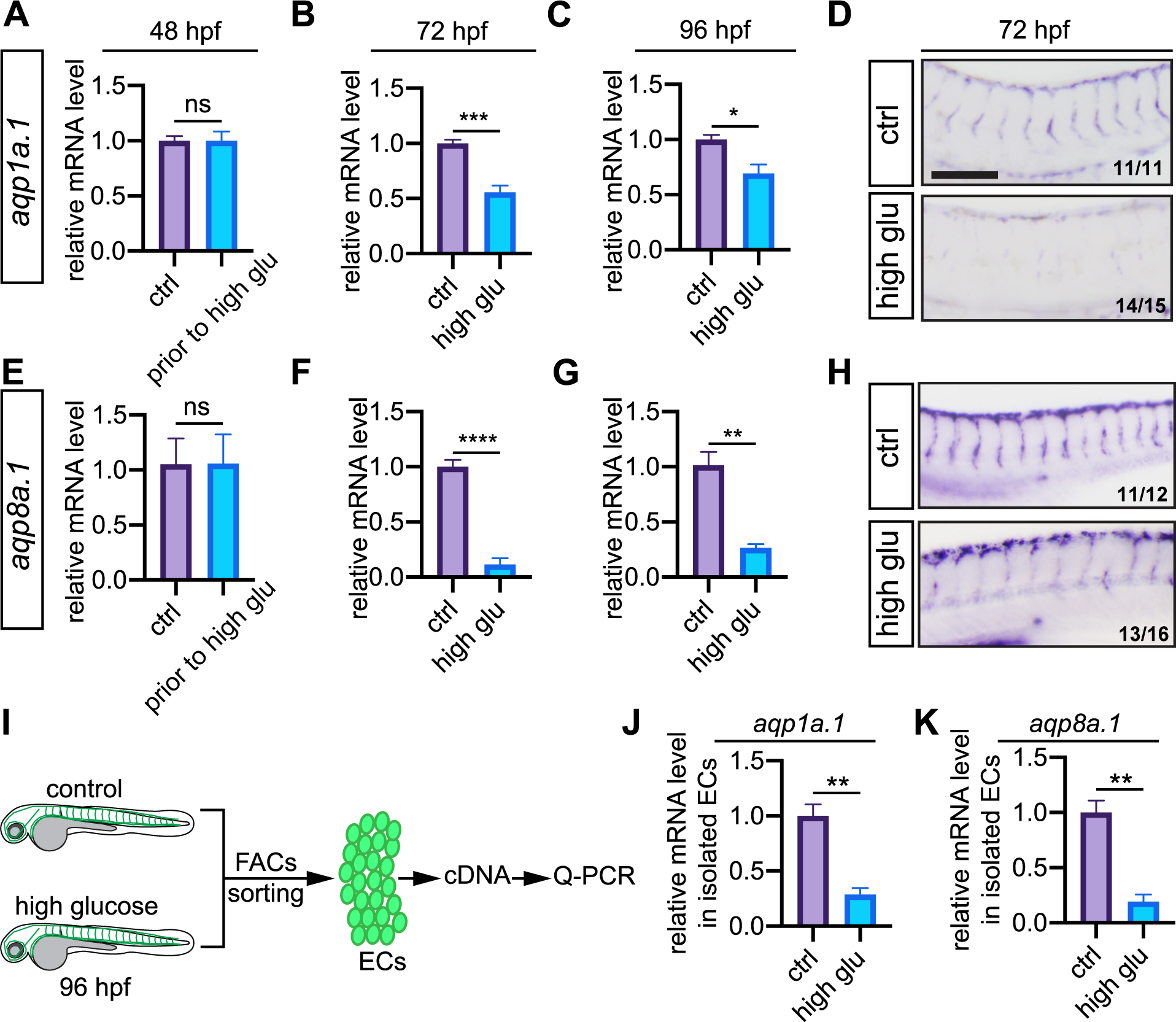
Zebrafish *aqp1a.1* and *aqp8a.1* are differentially expressed upon high glucose treatment. **A-H,** Expressions of *aqp1a.1* and *aqp8a.1* in zebrafish embryos are examined by Q-PCR and ISH. **A and E**, Expressions of *aqp1a.1* and *aqp8a.1* in zebrafish embryos (48 hpf) prior to high glucose treatment. **B and F**, Expressions of *aqp1a.1* and *aqp8a.1* in zebrafish embryos (72 hpf) after 24 hours high glucose treatment. **C and G**, Expressions of *aqp1a.1* and *aqp8a.1* in zebrafish embryos (96 hpf) after 48 hours high glucose treatment. A total of 10 embryos are utilized for RNA extraction, and 3 replicates are conducted. **D and H**, whole-mount *in situ* hybridized embryos showing the expressions of *aqp1a.1* and *aqp8a.1* in control and high glucose-treated embryos. The number in the bottom right corner of each panel is the number of embryos with typical phenotypes of the total observed embryos. **I,** Schematic representation of isolation of ECs from control and high glucose-treated *Tg(kdrl:EGFP)* zebrafish embryos for RNA extraction and Q-PCR. **J and K,** Quantification of *aqp1a.1* and *aqp8a.1* expressions in isolated ECs. Data are shown as mean ± s.e.m. A two-tailed, unpaired Student’s t-test is applied. ns, not significant. *, p<0.05. **, p < 0.01. ***, p < 0.001. ****, p < 0.0001. Scale bars, 100 μm.

### *aqp1a.1*/*aqp8a.1* loss-of-function reduce blood vessel diameters and gain-of-function increase

To gain an insight into the functions of EC-expressed AQPs in the vascular system, we performed gene functional analysis using a zebrafish model. We first used CRISPR/Cas9 genome editing method to knock out either *aqp1a.1* or *aqp8a.1* in *Tg(fli1ep:EGFP-CAAX)^ntu^*^666^ transgenic zebrafish (Fig.S6 and S7). Homozygous *aqp1a.1*-knockout mutants displayed reduced lumen size of the small intersegmental vessels (ISVs), which were generated from angiogenesis. However, the diameters of large vessels, including dorsal aortae (DA) and posterior cardinal vein (PCV), which are generated through vasculogenesis, were not significantly affected (Fig.5A). Upon *apq1a.1* gene deletion, the diameters of DV and PCV only decreased by 4.2% and 4.7% at 48 hpf, respectively, whereas the average diameter of ISVs showed an approximately 20% decrease (Fig.5B).

**Figure 5.**
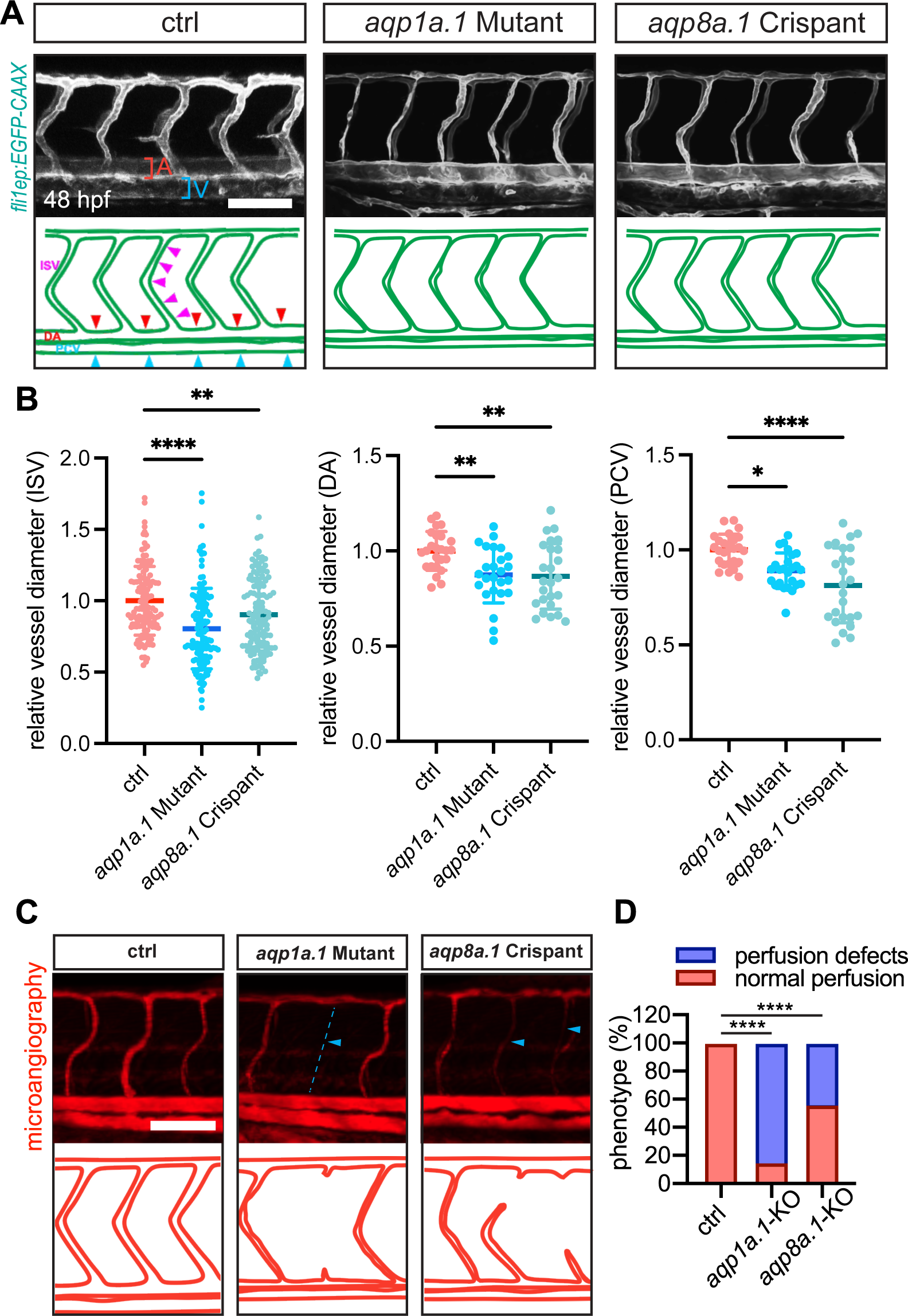
Knock-out of EC-enriched aquaporins (*aqp1a.1* or *aqp8a.1*) leads to the reduction of vascular diameter in zebrafish. **A,** Confocal images of ISV phenotypes in 48-hpf *Tg(fli1ep:EGFP-CAAX)^ntu^*^666^ control embryos (*n=18*), *aqp1a.1*-knockout embryos (*n=18*), and *aqp8a.1* crispants (*n=30*). Red and blue square brackets indicate artery and vein, respectively. Magenta, red and blue arrowheads in the representative sketch indicate the positions for vessel diameter measurement for ISV, DA, and PCV, respectively. **B,** Quantification of the diameters of ISV, DA and PCV. Each data point represents an individual vessel diameter measurement of ISV, DA, and PCV. Five fishes are analyzed for each group. Data are shown as mean ± s.e.m. One-way ANOVA analysis is applied. **C,** Microangiography of the trunk region in 72-hpf *Tg(fli1ep:EGFP-CAAX)^ntu^*^666^ control embryos (*n=18*), *aqp1a.1*-knockout embryos (*n=18*), and *aqp8a.1* crispants (*n=30*). Blue arrowheads indicate the perfusion-deficient ISVs. **D,** The incidence of normal and defective perfused ISVs in 72-hpf *Tg(fli1ep:EGFP-CAAX)^ntu^*^666^ control embryos (*n=18*), *aqp1a.1*-knockout embryos (*n=18*), and *aqp8a.1* crispants (*n=30*). One-way ANOVA analysis is applied. *, p<0.05. **, p < 0.01. ****, p < 0.0001. Scale bars, 100 μm.

To examine whether the vascular defects were caused by off-target effects of the CRISPR system, we knocked down *aqp1a.1* expression by injecting the *aqp1a.1* splicing morpholino into *Tg(fli1ep:EGFP-CAAX)^ntu^*^666^ embryos. The vascular defects were recapitulated and even enhanced within *aqp1a.1*-MO-injected zebrafish embryos (Fig.S8 and S9). These results suggested that Aqp1a.1 exerted a role in the regulation of vascular diameters. However, the homozygous or even heterozygous *aqp8a.1*-knockout mutants have not been obtained since all harvested embryos were genotyped as multiples of 3 base pairs deletion (Fig.S7). We were of the opinion that targeted deletion of the *aqp8a.1* gene resulted in embryonic lethality in zebrafish. Although it was hard to get *aqp8a.1*-knokout mutant, the G0 generation chimeras (*aqp8a.1* crispants) contained the *aqp8a.1*-knockout cells also displayed narrowed vessel lumens (Fig.5A). Compared with those of control, the diameters of DA, PCV and ISVs in *aqp8a.1*-KO G0 mutants decreased 11.2%, 9.8%, and 13.3%, respectively (Fig.5B). In order to test whether the vascular phenotypic change resulted from *aqp8a.1* loss of function, we injected *aqp8a.1*-specific morpholino into *Tg(fli1ep:EGFP-CAAX)^ntu^*^666^ embryos to block *aqp8a.1* pre-RNA splicing. The morpholino-mediated *aqp8a.1* knock-down led to reduced lumen diameters of blood vessels, which mimicked the phenotype caused by *aqp1a.1* loss-of-function (Fig.S8 and S9). Also, non-perfused or perfusion-defective ISVs were more frequently observed in *aqp1a.1* or *aqp8a.1* knock-out embryo based on microangiography analysis (Fig.5C and D). All the results suggested that loss-of-function of either *aqp1a.1* or *aqp8a.1* caused the narrowing of vascular diameters.

Next, we performed Aqp1a.1/Aqp8a.1 gain of function analysis to further confirm the contribution of EC-expressed aquaporins on blood vessel calibers. *Tg(fli1ep:aqp1a.1-mCherry)^ntu^*^667^ and *Tg(fli1ep:aqp8a.1-EGFP)^ntu^*^668^ lines were constructed to overexpress fluorescently-labeled Aqp1a.1 and Aqp8a.1 in zebrafish, respectively (Fig.S1B and C). Overexpressing either of them induced the enlargement of the blood vessel lumens (Fig.6A). Overexpression of *aqp1a.1* led to increases in DA and PCV diameter by 15.2% and 17.3%, respectively (Fig.6B and C), while the diameter of ISVs increased almost half of the control (approx. 47%) upon *aqp1a.1* overexpression (Fig.6D). The morphology of blood vessels in *aqp8a.1* gain of function embryos resembled the vascular phenotype of *aqp1a.1* overexpression (Fig.6A). Diameters of DA, PCV, and ISVs increased 22.4%, 23.2%, and 32.2% in *aqp8a.1* overexpressed zebrafish embryos, respectively (Fig.6B-D). Moreover, co-overexpression of *aqp1a.1* and *aqp8a.1* in zebrafish endothelial cells resulted in further dilation of DA, but the diameters of PCV and ISVs did not present a significant increase compared to single aquaporin (*aqp1a.1* or *aqp8a.1*) Overexpression (Fig.6A). The diameters of DA, PCV and ISVs in *aqp1a.1* and *aqp8a.1* co-overexpressed line exhibited 30.9%, 19.8%, and 39.0% elevation compared to the control (Fig.6B-D). These results suggested the role of Aqp1a.1 and Aqp8a.1 in promoting vascular lumen formation.

**Figure 6.**
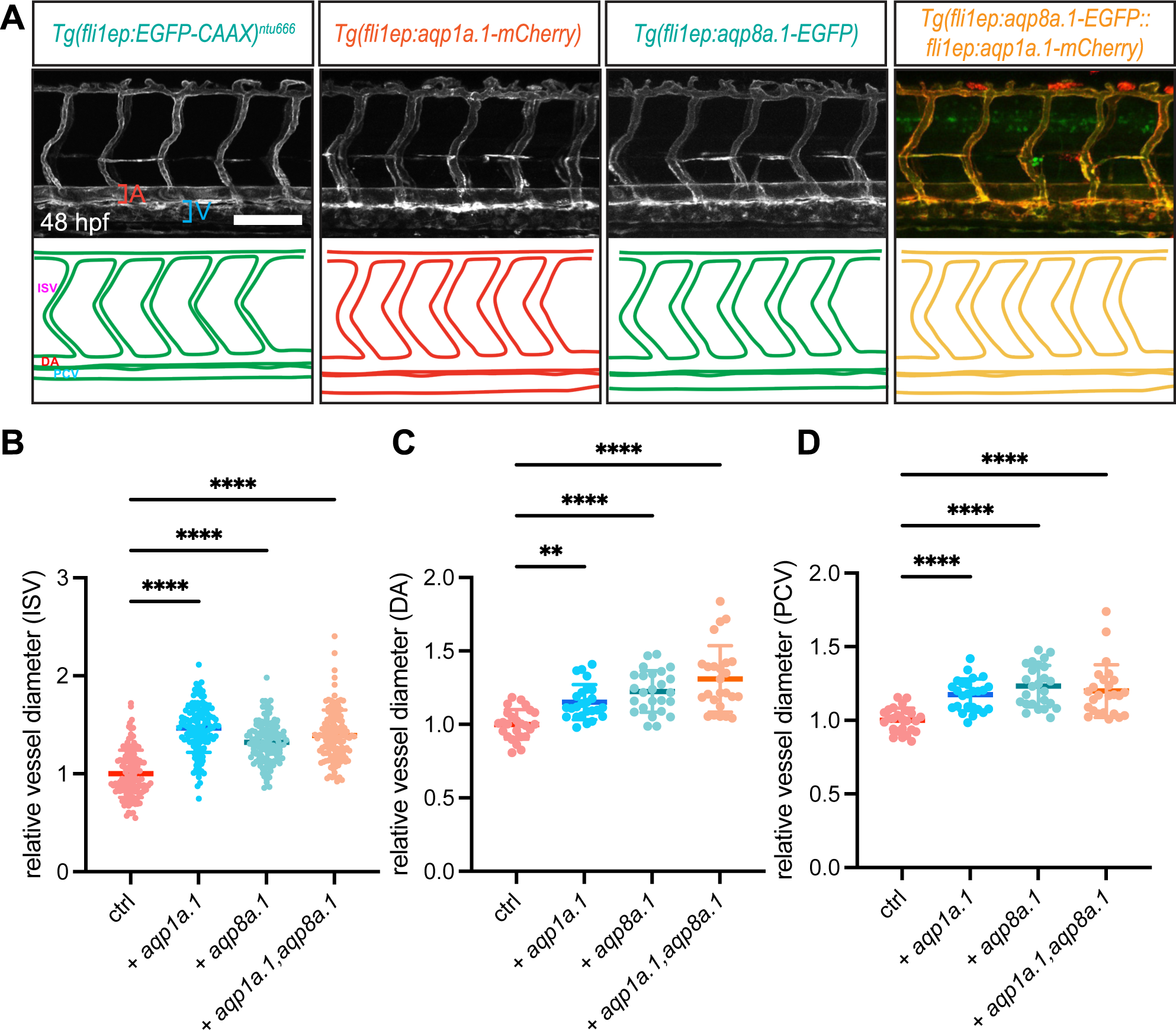
Overexpression of EC-enriched aquaporins (*aqp1a.1* or *aqp8a.1*) in ECs results in vessel lumen enlargement in zebrafish. **A,** Confocal images of ISV phenotypes in 48-hpf *Tg(fli1ep:EGFP-CAAX)^ntu^*^666^ transgenic line (control, *n= 18*), *Tg(fli1ep:aqp1a.1-mCherry)^ntu^*^667^ transgenic line (*aqp1a.1* overexpression, *n=25*), *Tg(fli1ep:aqp8a.1-EGFP)^ntu^*^668^ transgenic line (*aqp8a.1* overexpression, *n=23*), and *Tg(fli1ep:aqp8a.1-EGFP::fli1ep:aqp1a.1-mCherry)* double transgenic line (*aqp1a.1* and *aqp8a.1* co-overexpression, *n=25*). Red and blue square brackets indicate artery and vein, respectively. **B-D,** Quantification of the diameters of ISV, DA, and PCV. Each data point represents an individual vessel diameter measurement of ISV, DA, and PCV. Five fishes are analyzed for each group. Data are shown as mean ± s.e.m. One-way ANOVA analysis is applied. **, p < 0.01. ****, p < 0.0001. Scale bar, 100 μm.

In addition, we applied HgCl_2_ treatment on *Tg(fli1ep:EGFP-CAAX)^ntu^*^666^ zebrafish embryo to inhibit aquaporin function. Mercury has been commonly used to specifically block the water-channel activity of Aqp1a.1 and Aqp8a.1^29–32^. To screen the proper dosage for the treatment, we tried varied doses of HgCl_2_. It was shown that the embryos exposed to 1.6 μM HgCl_2_ from 22 to 48 hpf had no obvious morphological defects. Here, a lower concentration (0.4 μM) of HgCl_2_ was sufficient to narrow the diameter of the blood vessels (Fig.S10). Under this concentration, there were 15.4%, 15.7%, and 16.9% decrease in the diameters of DV, PCV, and ISVs, respectively. With a further increase in HgCl_2_ concentration, the diameters of all the aforementioned blood vessels reduced more remarkably (Fig.S10). As a result, blocking the channel activity of aquaporins led to defective vascular tube formation.

### Overexpression of *aqp1a.1* rescues hyperglycemia-induced decrease in vascular diameter

To elucidate whether the narrowed vessels induced by hyperglycemia is AQP-dependent, we establish a transgenic construct in which *apq1a.1* was chimerically upregulated in ECs. As shown in Fig.7A, the *pTol2-fli1ep-aqp1a.1-mCherry* plasmid, and Tol2 transposase mRNA were co-injected into 4-cell stage embryos of *Tg(fli1ep:EGFP-CAAX)^ntu^*^666^ progeny. By using this strategy, the mCherry-labeled Aqp1a.1 was mosaically expressed in the zebrafish vascular system. After injection, the embryos were first selected for mosaic mCherry fluorescence and subsequently immersed into glucose solution from 48 hpf for 24 h prior to confocal imaging analysis (Fig.7B). As expected, a few ISVs exhibited both mCherry and GFP expression, and these ISVs displayed wider lumens than the ones without mCherry expression (Fig.7C and D). These results demonstrated that Overexpression of *aqp1a.1* recovered the defective lumen formation caused by hyperglycemia.

**Figure 7.**
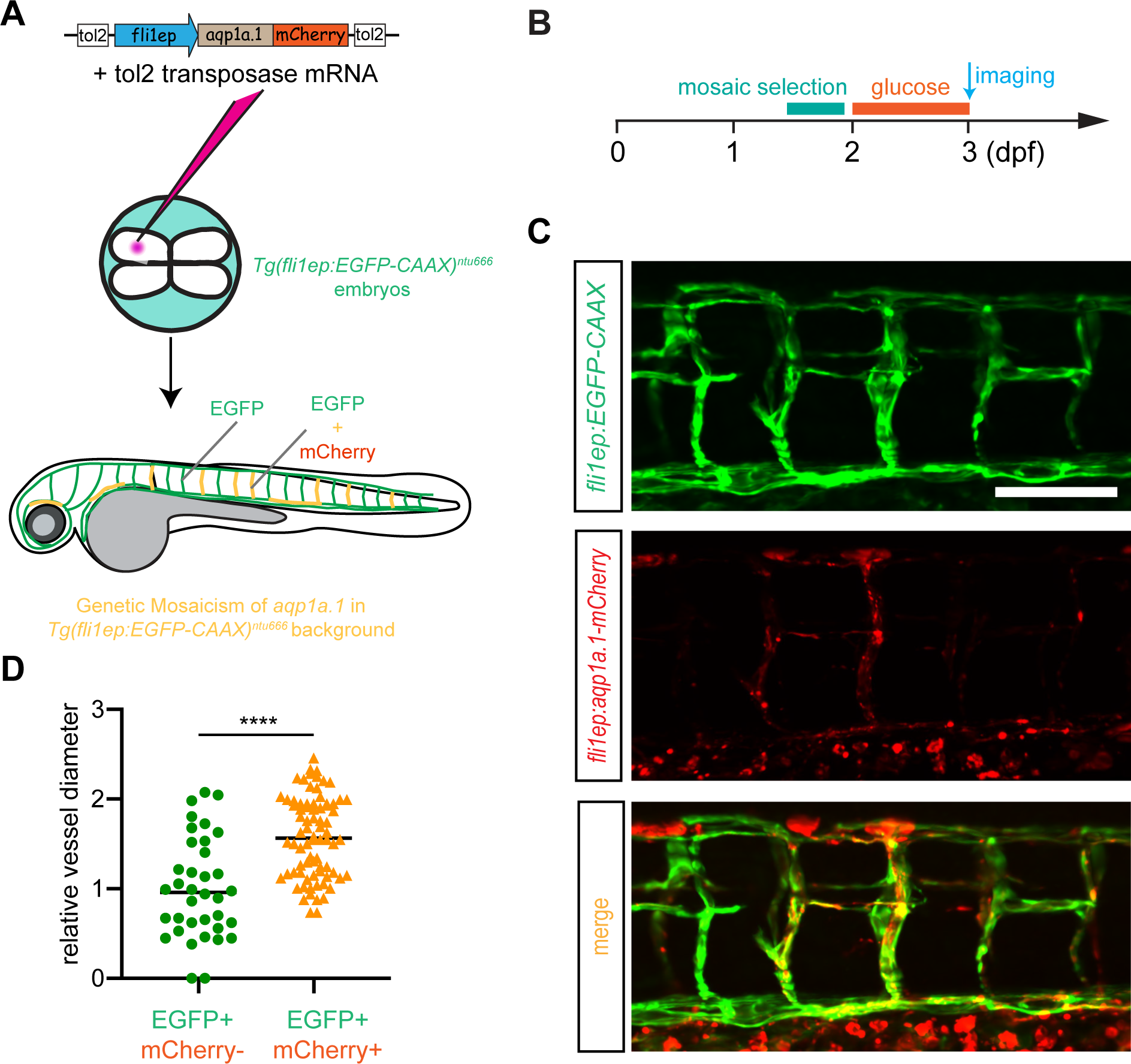
Overexpression of zebrafish *aqp1a.1* rescues hyperglycemia-caused narrowing of vascular diameter. **A,** Schematic representation of establishment of *aqp1a.1*-overexpressed transgenic line under *Tg(fli1ep:EGFP-CAAX)^ntu^*^666^ background. DNA construct and Tol2 transposase mRNA are co-injected at the 4-cell stage to establish a chimera line, in which the mCherry-labeled Aqp1a.1 is mosaically expressed in the vascular system. **B,** Embryos are selected and immersed into high glucose solution from 2 to 3 dpf prior to imaging. **C,** Representative confocal images of ISVs in 3-dpf *Tg(fli1ep:EGFP-CAAX::fli1ep:apq1.1-mCherry)* double transgenic line (*n=40*). **D,** Quantification of the lumen diameters in ISVs with or without Aqp1a.1 expression. Each data point represents an individual vessel diameter measurement of ISV, DA, and PCV. Five fishes are analyzed for each group. Data are shown as mean ± s.e.m. A two-tailed, unpaired Student’s t-test is applied. ****, p < 0.0001. Scale bar, 100 μm.

### AQP1 is required for vascular tube formation but not for EC differentiation

As the zebrafish *aqp1a.1* homologue, human *AQP1* is specifically and strongly expressed in most microvascular endothelial cells ^14^. However, human *AQP8* (zebrafish *aqp8a.1* homologue) is mainly presented in epithelial cells of varied tissues but barely detected in endothelial cells ^33^. Our results also demonstrated that *AQP1* was specifically expressed in ECs (including HUVECs and isolated human ECs), similar to the maker genes *CD31* and *CD144*, but *AQP8* expression was almost undetectable in ECs (Fig.S11A and B). In comparison to ECs, *AQP1* and *AQP8* were seldom expressed in human ESCs (H1 line) (Fig.S11D). However, H1-derived ECs (H1-ECs) (from day 4 onwards) showed an increased expression of *AQP1,* but the *AQP8* expression was not altered (Fig.S11C and D).

To further examine the role of AQP1 in vessel formation, we generated two *AQP1*-knockout human ESC lines with H1 cells (Δ*AQP1*-1H1 and Δ*AQP1*-2H1) through a previously established episomal vector-based CRISPR/Cas9 (epiCRISPR) system (Fig.8A) ^26^. The embryoid body (EB)-based strategy for EC differentiation was utilized with wild-type H1 cells (WT-H1) and those two AQP-knockout lines. ECs arise from WT-H1 (WT-H1-ECs), Δ*AQP1*-1H1 (Δ*AQP1*-1H1-ECs), and Δ*AQP1*-2H2 (Δ*AQP1*-1H2-ECs) were respectively subjected to CD31 and CD144 immunostaining followed by FACS analysis. Our results showed that there was no significant difference in the percentage of CD31+ and CD144+ populations among these three EC groups differentiated from WT and AQP1-knockout lines (Fig.8B-D). Quantitative PCR analysis confirmed the loss of *AQP1* in Δ*AQP1*-1H1-ECs and Δ*AQP1*-1H2-ECs, but the expressions of maker genes *CD31* and *CD144* were comparable in these three derived ECs (Fig.8E-G). Although these three types of H1-ECs were all capable of forming angiogenic sprouts when seeded onto Matrigel, the formation of tube-like structures was severely inhibited, and the number of junctions and the total length of the sprouts significantly declined in *AQP1*-deleted cells (Fig.8H-M). All these findings suggested that endothelial AQP1 is a novel regulator in vascular tube formation but had no effects on EC differentiation.

**Figure 8.**
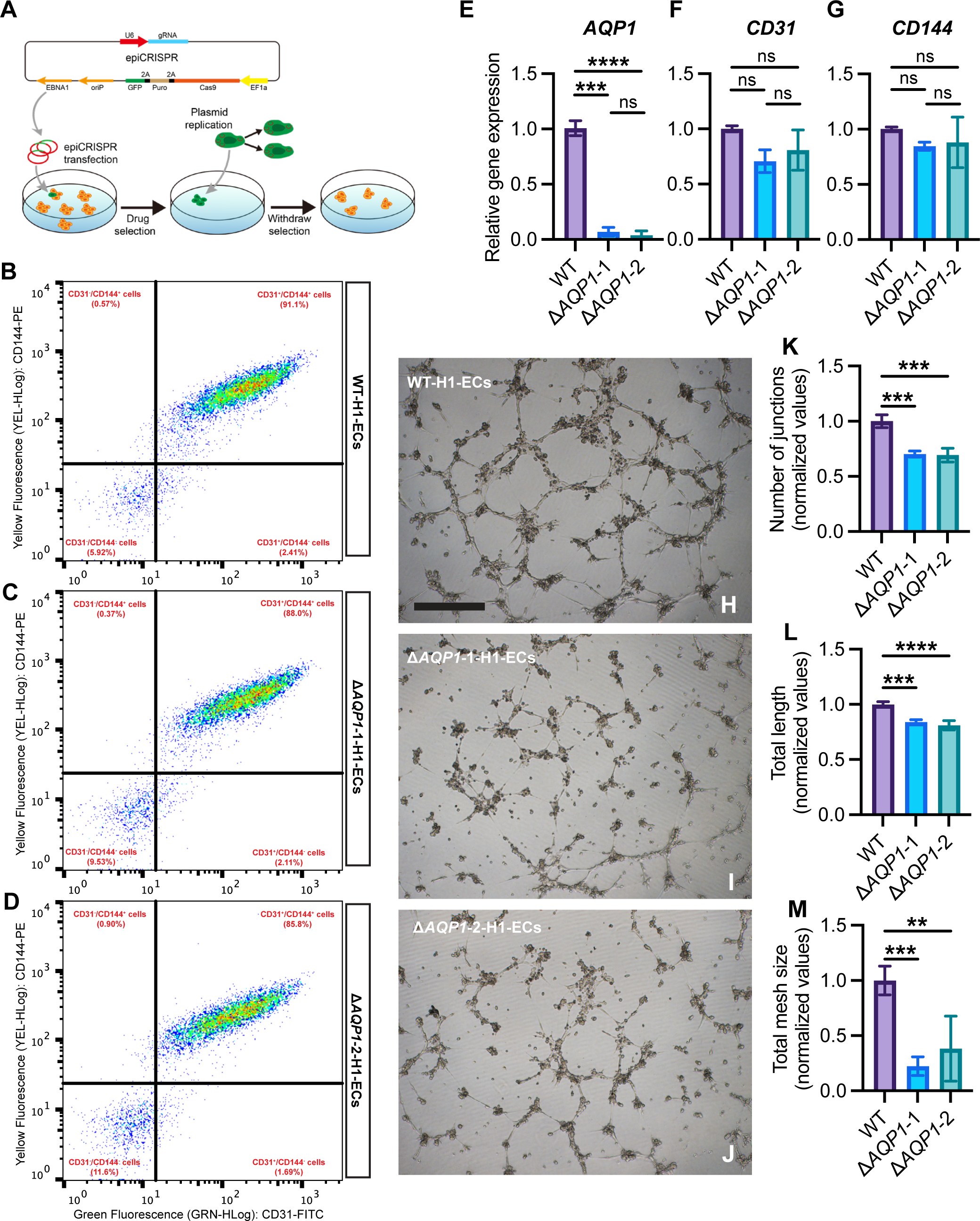
*AQP1* deletion does not affect EC differentiation from H1 stem cells but inhibits tube formation of the derived ECs. **A,** Schematic representation of the epiCRISPR system for *AQP1* gene knock-out in H1 cell line. The vector contains a U6 promoter-driven gRNA scaffold, an EF1a promoter-driven Cas9 fused to puromycin resistance gene and GFP with P2A peptides, and OriP/EBNA1 elements for the plasmid replication in eukaryotes. The epiCRISPR vector can replicate in H1 cells and can be partitioned into daughter cells. **B-D,** Flow cytometry analyses showing the percentages of cells that express CD31 and CD144 endothelial markers in ECs derived from H1 cell line (**B**) and two *AQP1*-knockout H1 cell lines, Δ*AQP1*-1-H1 (**C**) and Δ*AQP1*-2-H1 (**D**). **E-G,** Q-PCR analyses confirming the gene expressions of *AQP1*, *CD31,* and *CD144* in ECs derived from H1 (**E**), Δ*AQP1*-1-H1 (**F**), and Δ*AQP1*-2-H1 (**G**) cell lines. **H-J,** Representative microscopy images showing the tube formation of ECs derived from H1 cell line (**H**), Δ*AQP1*-1-H1 (**I**), and Δ*AQP1*-2-H1 (**J**) cell lines. The derived ECs are seeded on the matrigel for 6 hours before being subjected to analysis. The number of junctions, the total length of tubes, and total mesh size of the vascular network formed by ECs derived from H1 (*n=6*), Δ*AQP1*-1-H1 (*n=6*), and Δ*AQP1*-2-H1 (*n=6*) cells are shown in **K, L, and M**, respectively. Data are shown as mean ± s.e.m. One-way ANOVA analysis is applied. ns, not significant. *, p<0.05. **, p < 0.01. ***, p < 0.001. ****, p < 0.0001. Scale bars, 100 μm.

### AQPs regulate vascular diameter through intracellular vacuoles

Given that intracellular vacuole fusion is an essential way of vascular lumenization in zebrafish ISVs ^7^, whether AQPs involve in this process is undetermined. By expressing the fluorescently-labeled *aqp1a.1* and *aqp8a.1* in human umbilical vein endothelial cells (HUVECs), we found that both Aqps were co-localized on the surface of intracellular vacuoles in ECs (Fig.9A-C). Co-Injection Tol2 transposase with pDest-fli1a-aqp8a.1-EGFP or pDest-fli1a-aqp1a.1-mCherry expression vector into one-cell stage zebrafish embryos revealed that both Aqps were expressed and highly enriched in the subcellular vacuoles of zebrafish ISVs (Fig.9D). Moreover, Aqp1a.1 was co-localized with Cdc42, a marker for intracytoplasmic vesicles ^7, 34^, in sprouting ISVs of double transgenic line *Tg(fli1:EGFP-cdc42wt::fli1ep:apq1.1-mCherry)* (Fig.9E). Meanwhile, *aqp1a.1* knock-out in combination with *aqp8a.1* knock-down led to the reduction of intracellular vacuoles and defective endothelial tube assembly in ISVs of the *Tg(fli1:EGFP-cdc42wt)* transgenic line (Fig.9F and G). In addition, the total size of endothelial vacuoles significantly declined upon loss of Aqps in zebrafish ISVs (Fig.9H). Next, we examined the effects of the loss of AQP1 on the formation of intracellular vacuoles between H1-derived ECs and AQP1 knock-out H1 cell line-derived ECs *in vitro*. Compared to H1-derived ECs (WT-H1-ECs), the loss of AQP1 led to a significant decrease in the total vacuole size of ECs (Fig.9I and J). All these results indicated that endothelial AQPs displayed cytoplasmic vacuolar localization and were required for intracellular vacuole-mediated endothelial tube formation.

**Figure 9.**
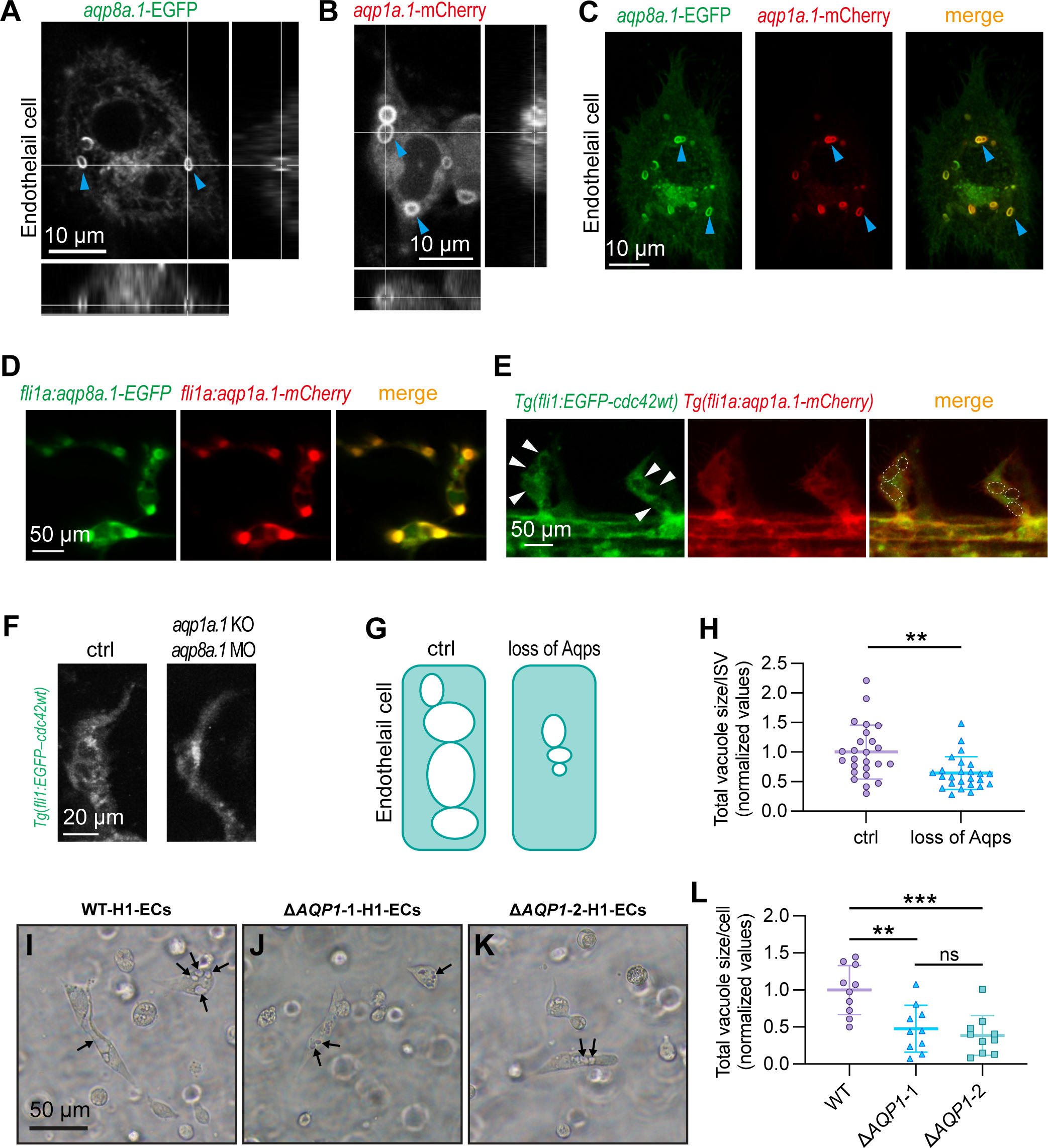
Endothelial AQP facilitates intracellular vacuole formation in ECs *in vitro* and *in vivo*. **A-C**, Fluorescently-labeled Aqp1a.1 and Aqp8a.1 are expressed on the surface of cytoplasmic vacuoles (blue arrowheads) in HUVECs. Subpanels at the bottom and right of **A and B** are cross-sections of the intracellular vacuoles. **C,** Aqp1a.1 and Aqp8a.1 co-localize at the intracellular vacuoles. Scale bars, 10 μm. **D,** EGFP-labeled Aqp1a.1 and mCherry-labeled Aqp8a.1 are expressed and co-localized at the intracellular vacuoles in zebrafish ISVs. **E,** Confocal imaging of zebrafish ISVs in *Tg(fli1:EGFP-cdc42wt::fli1a:aqp1a.1-mCherry)* double transgenic line, in which the EGFP-labeled Cdc42 protein marks the vacuolar structures (white arrowheads), and Aqp1a.1 is labeled by mCherry. Aqp1a.1 co-localizes with intracellular vacuoles (dashed circles) (*n=20*). Scale bars, 50 μm. **F,** Knock-out *aqp1a.1* and knock-down *aqp8a.1* in zebrafish lead to defective vascular lumen formation and reduced intracellular vacuoles in ISVs (*n=19*). Scale bars, 20 μm. **G,** Schematic representation of intracellular vacuole changes upon loss of Aqps. **H,** Quantification of the total size of intracellular vacuoles in zebrafish ISVs. Each point represents the total size of intracellular vacuoles in an individual ISV. Five fishes are analyzed for each group. Data are shown as mean ± s.e.m. A two-tailed, unpaired Student’s t-test is applied. **, p < 0.01. **I-J,** Representative images of intracellular vacuoles (arrows) in ECs derived H1 cell line (**I**), Δ*AQP1*-1-H1 (**J**) and Δ*AQP1*-2-H1 (**K**) cell lines. Scale bars, 50 μm. **L,** Quantification of the total size of intracellular vacuoles in cultured ECs. Each point represents the size of intracellular vacuoles per cell. One-way ANOVA analysis is applied. ns, not significant. **, p < 0.01. ***, p < 0.001.

## Discussion

Diabetic retinopathy (DR) is a common vascular complication of diabetes and also a leading cause of vision loss among working-age populations ^35^. It is often accompanied by a series of retinal vascular lesions, including reduced vessel perfusion, damaged vasculature, increased vascular permeability, and neovascularization ^4^. Hyperglycemia is an inextricable reason for DR, and the diameter changes in retinal vessels have been recognized as a clinical sign of DR^36, 37^. In our study, we observed a significant downregulation of *AQP1* in the retinas of patients with diabetes and high glucose-treated HRMECs. These observations indicate a novel pathophysiological link between AQP1 and DR. Aquaporins (AQPs) belongs to a superfamily of major intrinsic proteins (MIPs) and serves as channel proteins to transport water and/or small solutes across plasma membrane ^10, 11, 38^. Based on their ability to transport water or other solutes, such as glycerol, the aquaporin superfamily can be divided into two subfamilies: aquaporins and aquaglyceroporins. However, whether AQPs play roles in the progression of pathological vascular complications is poorly understood. Discoveries in vascular research have been limited by a shortage of a model for imaging the blood vessels at high resolution *in vivo*. To address this, the zebrafish model was utilized in our study, which offers superior properties in cardiovascular research. In zebrafish, 14 mammalian AQP homologues have been cloned, and their expression has been examined with WISH ^39^. The unique spatiotemporal distribution of individual aquaporins during zebrafish embryogenesis implied their distinct roles in embryonic development. Although aquaporins, particularly AQP1, have been reported to be strongly expressed in ECs and proliferating tumor microvessels in humans ^12, 14, 40, 41^, their functions in the vascular system remain to be elucidated. Among all these aquaporin homologues, zebrafish *aqp1a.1* and *aqp8a.1* were endothelium-enriched. To further investigate the effects of EC-expressed Aqp1a.1 and Aqp8a.1 on angiogenesis during zebrafish development, we examined the vascular phenotypic changes upon *aqp1a.1*/*aqp8a.1* loss-of-function and gain-of-function. Either *aqp1a.1* or *aqp8a.1* disruption in zebrafish embryos narrowed dorsal aorta, cardinal vein, and ISVs, which were more sensitive to AQPs loss of function. In contrast, overexpressing *aqp1a.1* or *aqp8a.1* led to increases in vascular diameters. Moreover, blockage of the function of AQPs by mercury resulted in the narrowing of the blood vessels as well. Taken together, these observations suggested that AQPs modulated the blood vessel diameters during both vasculogenesis and angiogenesis. In addition, both *aqp1a.1* and *apq8a.1* were localized in the intracellular vacuoles in cultured ECs as well as the ECs of sprouting ISVs, and loss of Aqps caused the reduction of those vacuoles. Since zebrafish ISVs adopted the intracellular vacuoles fusion as the main strategy for vascular lumenization, the EC-enriched Aqps might play a critical role in this biological process, supporting two previous notions that the vascular lumen formation was driven by the intercellular fusion of endothelial vacuoles ^7^ and AQP was expressed on cytoplasmic vesicles and required for lumen extension ^31^.

Zebrafish has been widely used in diabetic studies ^42–44^, and a hyperglycemic state and a series of diabetic vascular complications can be easily simulated by immersing the zebrafish embryos or adults in highly concentrated glucose solution ^45, 46^. In our data, high glucose treatment led to vascular defects, which were well-matched with the defective vascular phenotype in *aqp1a.1*/*aqp8a.1* loss-of-function. Under the hyperglycemic condition, the IOC in eyes and ISVs were narrowed significantly. Furthermore, we discovered that zebrafish *aqp1a.1* and *aqp8a.1* were downregulated upon high glucose exposure, which was consistent with our observation that *AQP1* was downregulated in human diabetic retinas and high glucose-treated HRMECs. Although elevated glucose was able to cause cardiac defects and dimmish blood flow ^46, 47^, which might disrupt vascular perfusion and lumenization, the vascular defects could be recovered to normal when overexpressing *aqp1a.1* (human *AQP1* ortholog) in ECs of zebrafish hyperglycemic model. All these results were in support of the viewpoint that the deficiency in blood vessel formation is not a secondary effect of a dysfunctional heart, and the EC-expressed aquaporins are correlated with hyperglycemia-related vascular complications. Sada et al. has ever mentioned that Overexpression of AQP1 attenuates hyperglycemia-induced vascular complications ^48^. In their proposed working model, hyperglycemia leads to high oxygen consumption, which promotes mitochondrial ROS production and induces cellular hypoxia. Afterward, the mitochondrial ROS and cellular hypoxia promote the overproduction of endothelin-1 and fibronectin and induce apoptosis, which further leads to diabetic vascular complications. Based on our data, the attenuated hypoxia might be due to an improvement of the vascular perfusion induced by the Overexpression of AQP1. Based on our observations, AQP1 may attenuate hypoxia under hyperglycemic conditions by improving vascular perfusion. Meanwhile, AQP1 can also suppress the hypoxic state by functioning as the oxygen channel in ECs ^49^ and then prevent the aforementioned hyperglycemia-induced vascular damage. In addition to AQP1, other non-EC-expressed aquaporins are also associated with diabetes. Zhang et al. have found that AQP11 was down-regulated in DR and resulted in intracellular edema in DR subsequently ^50^. AQP2 was also reported to be involved in diabetic kidney diseases ^51, 52^. However, to our knowledge, it was the first time that AQP1 was reported to be involved in physiological as well as pathological vascular lumen formation.

Our investigations on retina samples from diabetic patients and the zebrafish DM model emphasized that EC-enriched aquaporins are associated with vascular diameter regulation in hyperglycemia. Functional inhibition of aquaporins recapitulated vascular alterations of the high-glucose treated zebrafish model, and overexpression of aquaporin in ECs rescued the defect. In summary, the current study suggests that EC-enriched aquaporins have a role in developmental and pathological blood vessel lumenization, and they might be potential targets for gene therapy to cure diabetes-related vascular lumenization defects.

## Acknowledgments

This study was supported by grants from the National Natural Science Foundation of China (https://www.nsfc.gov.cn/; 81870359 and 2018YFA0801004 received by DL; 82000458 received by CC). The funders had no role in study design, data collection and analysis, the decision to publish, or the preparation of the manuscript.

## Author Contributions

DL and SH supervised and designed this project. CC, DL, and YQ wrote the manuscript. DL, CC, YQ, YX, and XS analyzed the data. CC, YQ, YX, XW, JG, WL, and YW performed the experiments; YX, LC, and LW collected the clinical data. DL, YW, and SH revised the manuscript.

## Supplementary Figure Legend

**Supplementary Figure 1.**
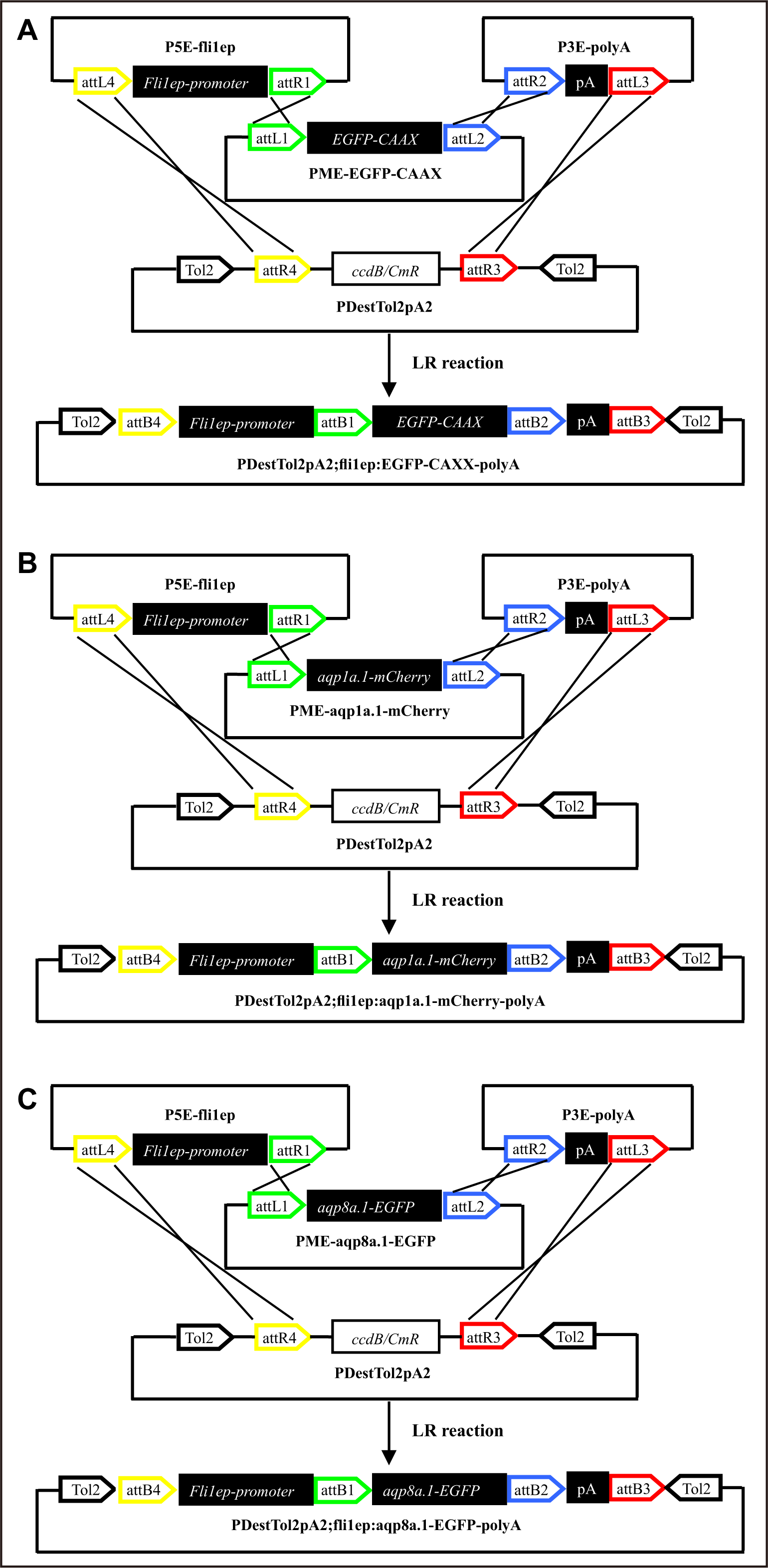
The diagram for showing the constructs of three transgenic lines.

**Supplementary Figure 2.**
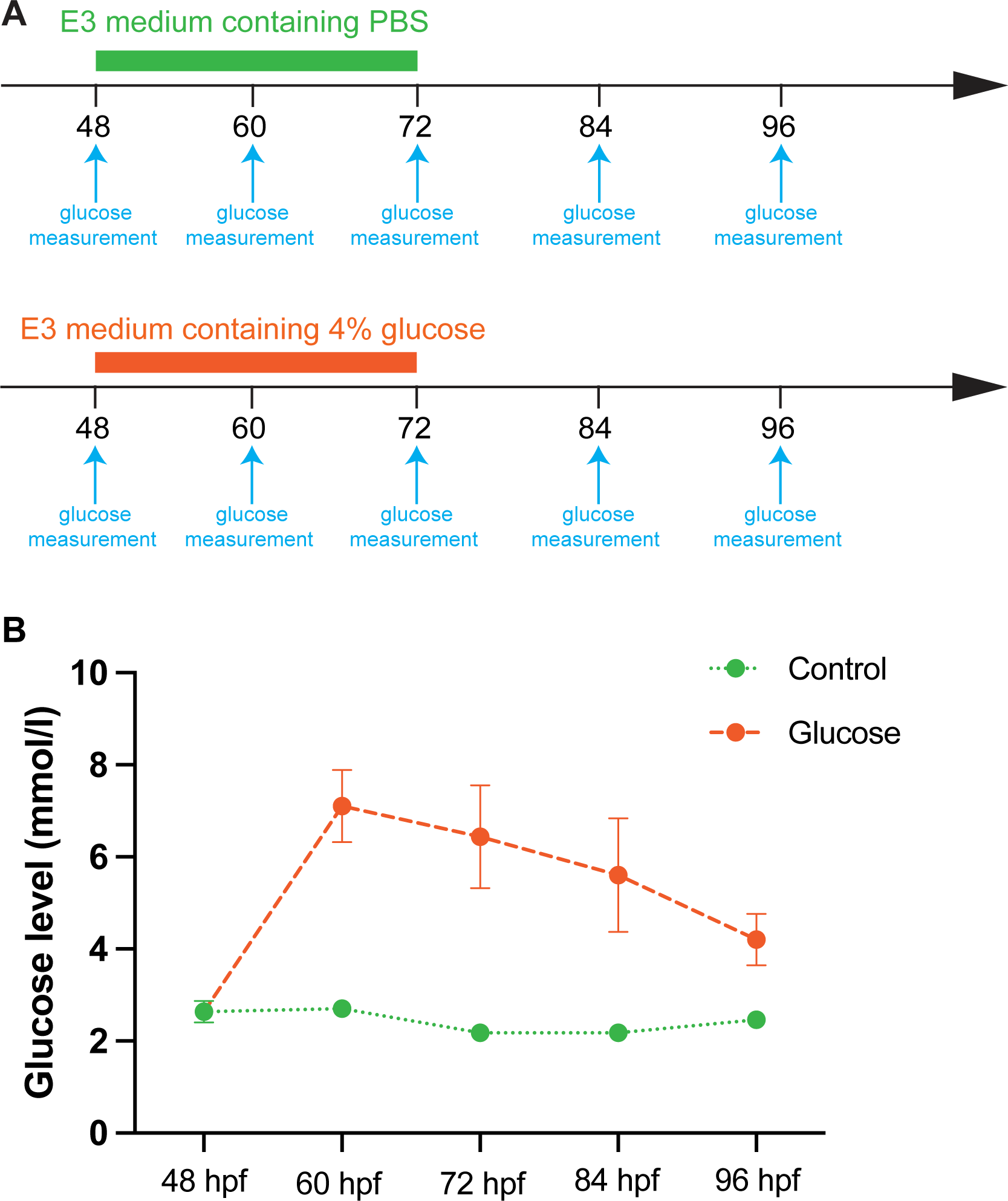
High glucose treatment of zebrafish embryos. **A,** The timeline of glucose treatment of zebrafish embryos. **B,** The glucose level of body fluid of zebrafish embryos.

**Supplementary Figure 3.**
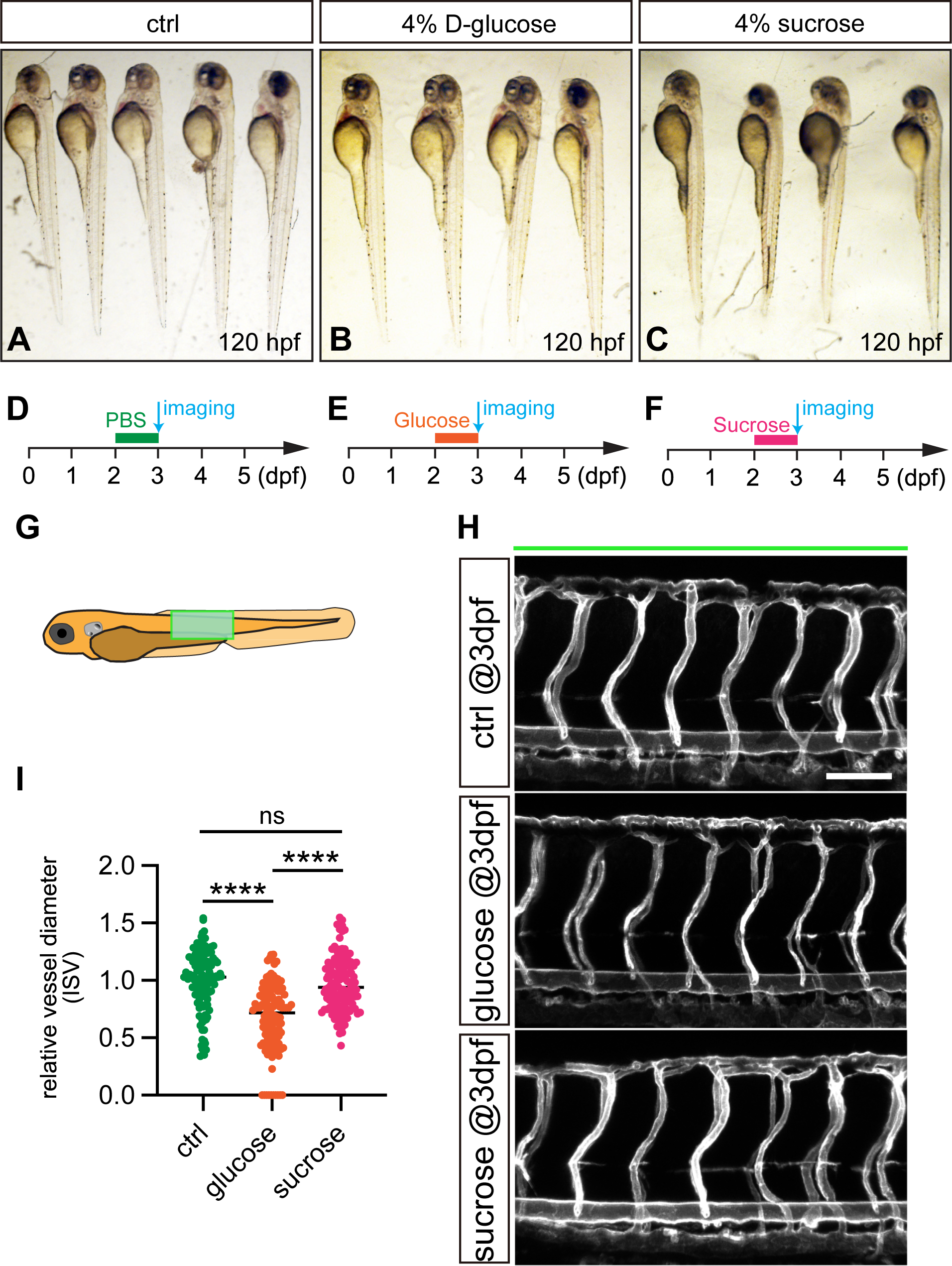
Comparative analysis of the effects of high glucose and high sucrose on vascular development in zebrafish embryos. **A-C,** Stereo microscopic analysis of control, glucose, and sucrose treated embryos in bright field. Zebrafish embryos are immersed into PBS buffer **(D)**, high glucose solution (4% D-glucose w/v) **(E)**, and high sucrose solution (4% sucrose w/v) **(F)** from 2 to 3 dpf. Confocal imaging of the green rectangle outlined area **(G)** is performed afterward. **H,** Representative confocal images of ISV phenotypes in control, glucose, and sucrose treated embryos. Scale bars, 100 μm. I, Quantification of ISV diameter. One-way ANOVA analysis is applied. ns, not significant. ****, p < 0.0001.

**Supplementary Figure 4.**
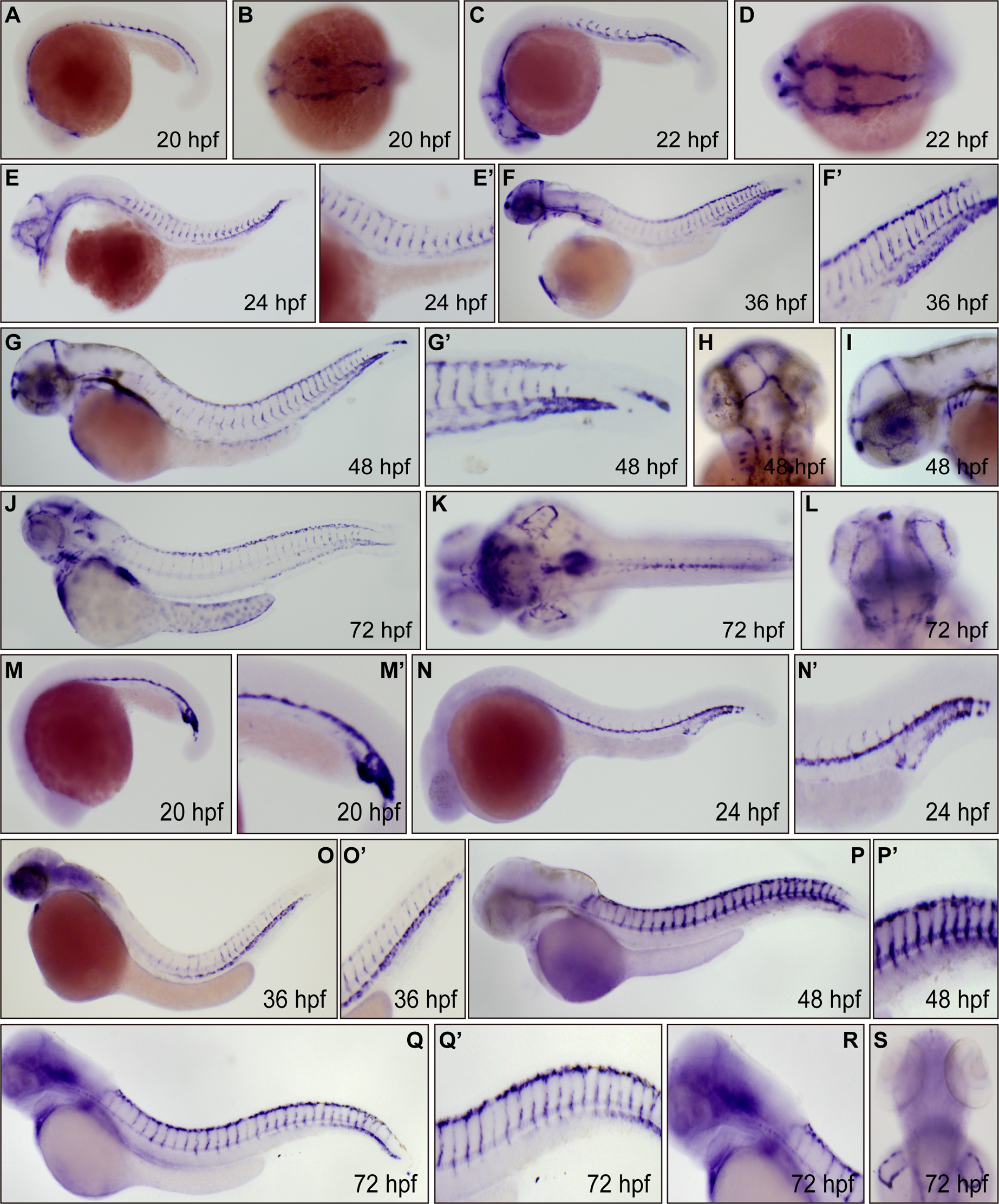
WISH analysis of *aqp1a.1* and *aqp8a.1*. **A-L,** The *aqp1a.1* expression in 20, 22, 24, 36, 48, and 72 hpf embryos. **M-S,** The *aqp8a.1* expression in 20, 24, 36, 48, and 72 hpf embryos.

**Supplementary Figure 5.**
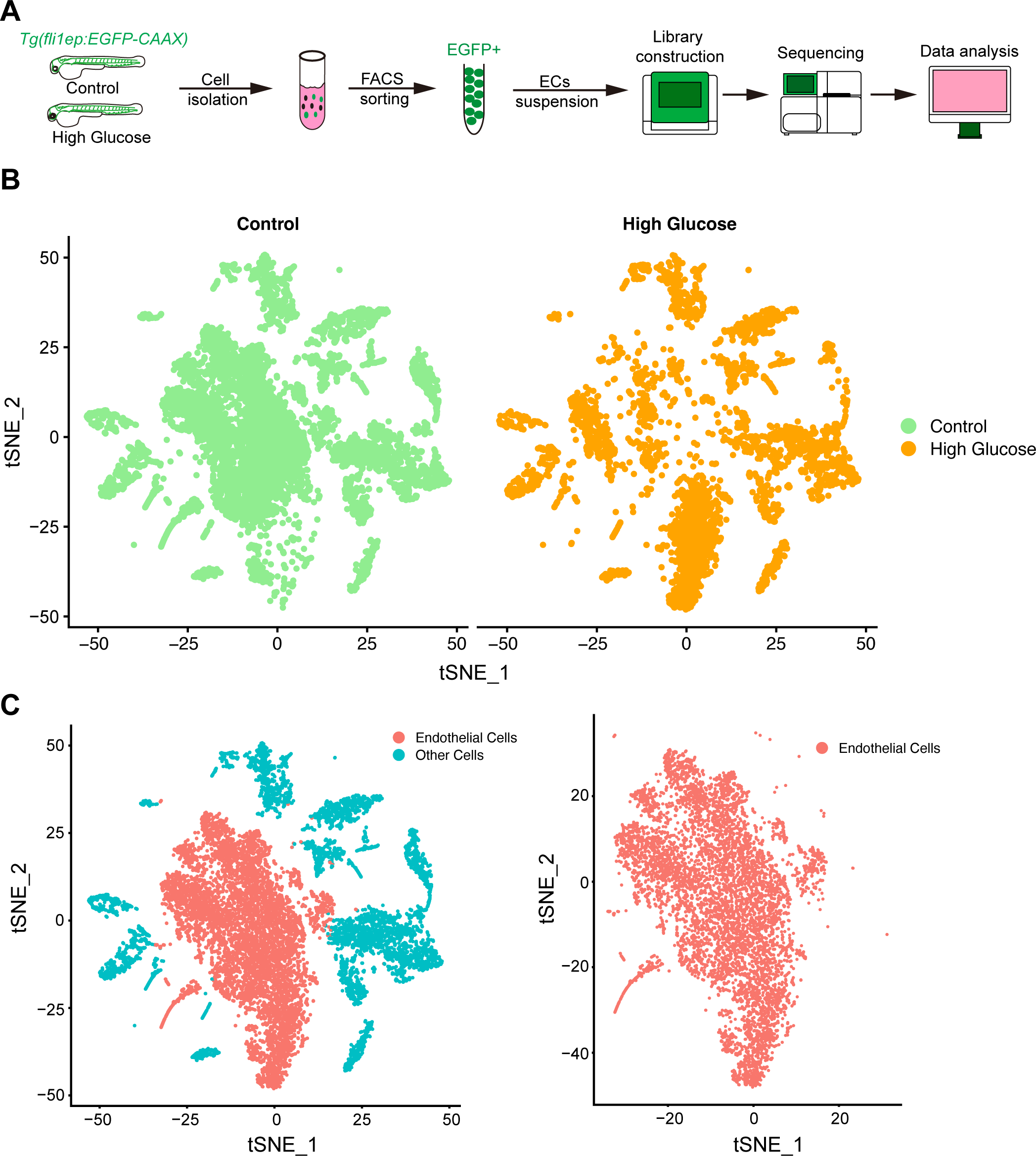
Single-cell RNA-seq profiling of GFP-positive cells from *Tg(fli1ep:EGFP-CAAX)^ntu^*^666^ zebrafish embryos. **(A)** Overview of the single-cell isolation and analysis workflow. **(B)** t-SNE plots of cells isolated from control and high glucose-treated groups. **(C)** Cells positive for *pecam1*, *cdh5*, and *kdrl* are separated.

**Supplementary Figure 6.**
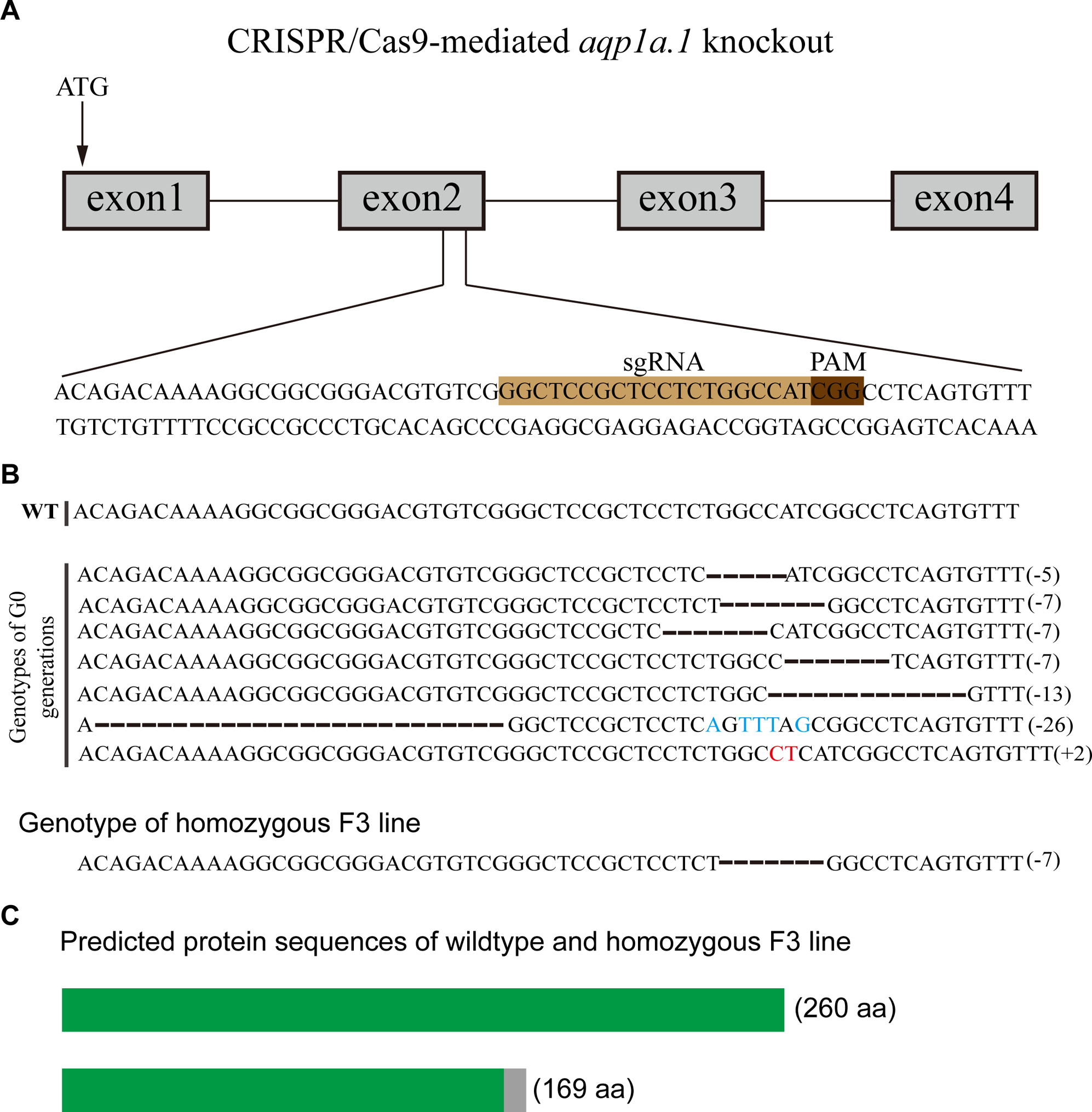
Generation of zebrafish *aqp1a.1* knock-out using CRISPR/Cas9. **A,** Schematic diagram showing gRNA targeting a site on the second exon of *aqp1a.1* gene. Starting codon (ATG) site is indicated by the arrow. **B,** Genotypes of G0 generations and homozygous F3 line. Numbers in the brackets show the number of nucleotides that were deleted (−) or inserted (+). An inserted nucleotide is in red. WT, wild-type. **C,** Schematic diagram showing the predicted proteins encoded by the wild-type and homozygous F3 line. The mutants are reading frameshift mutations that result in truncated proteins. The gray rectangles indicate the wrong-coded amino acid sequences.

**Supplementary Figure 7.**
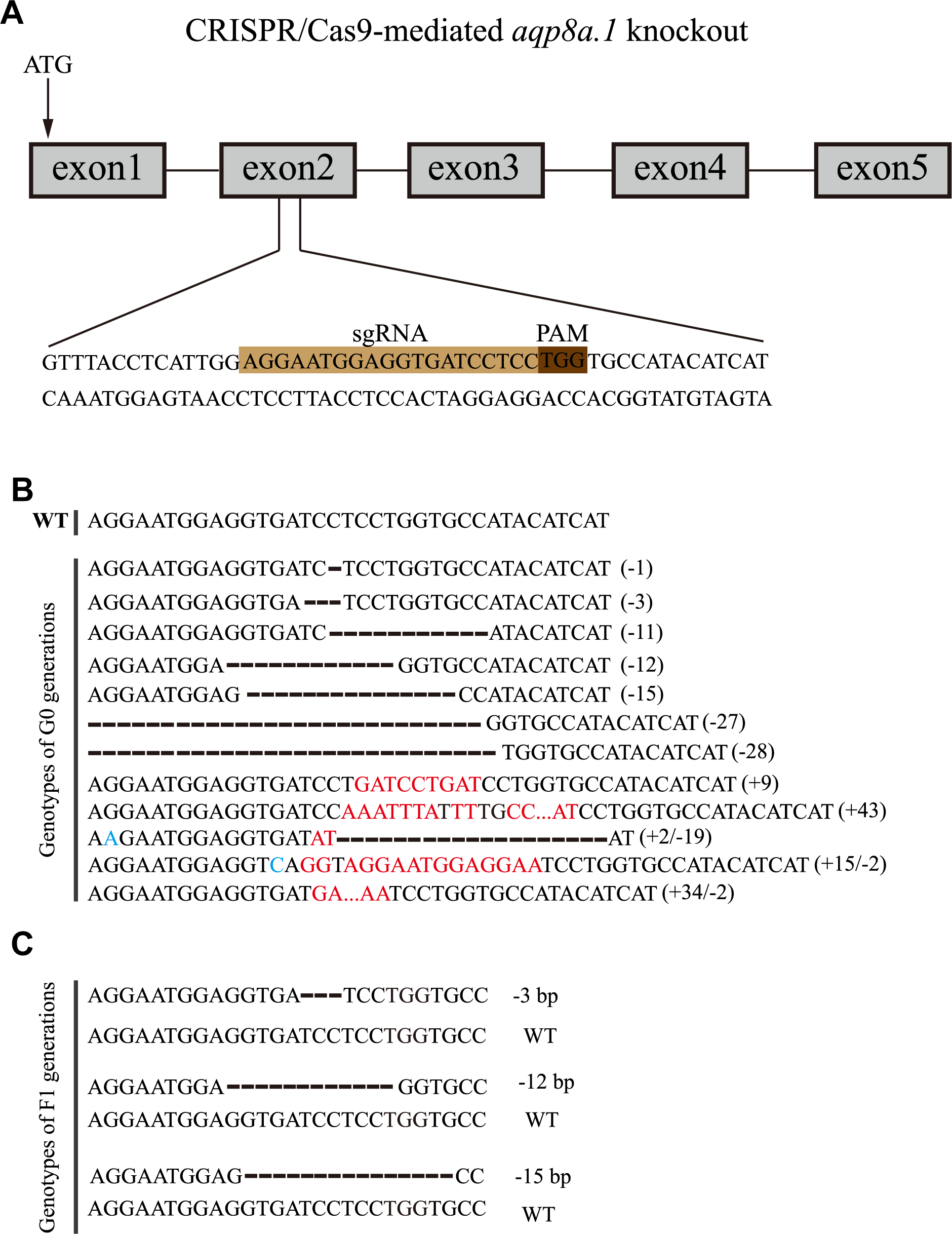
Generation of zebrafish *aqp8a.1* knock-out using CRISPR/Cas9. A. Schematic diagram showing gRNA targeting a site on the second exon of *aqp8a.1* gene. Starting codon (ATG) site is indicated by the arrow. B. Genotypes of G0 generations. Numbers in the brackets show the number of nucleotides that were deleted (−) or inserted (+). An inserted nucleotide is in red. C. Genotypes of heterozygous F1 generations.

**Supplementary Figure 8.**
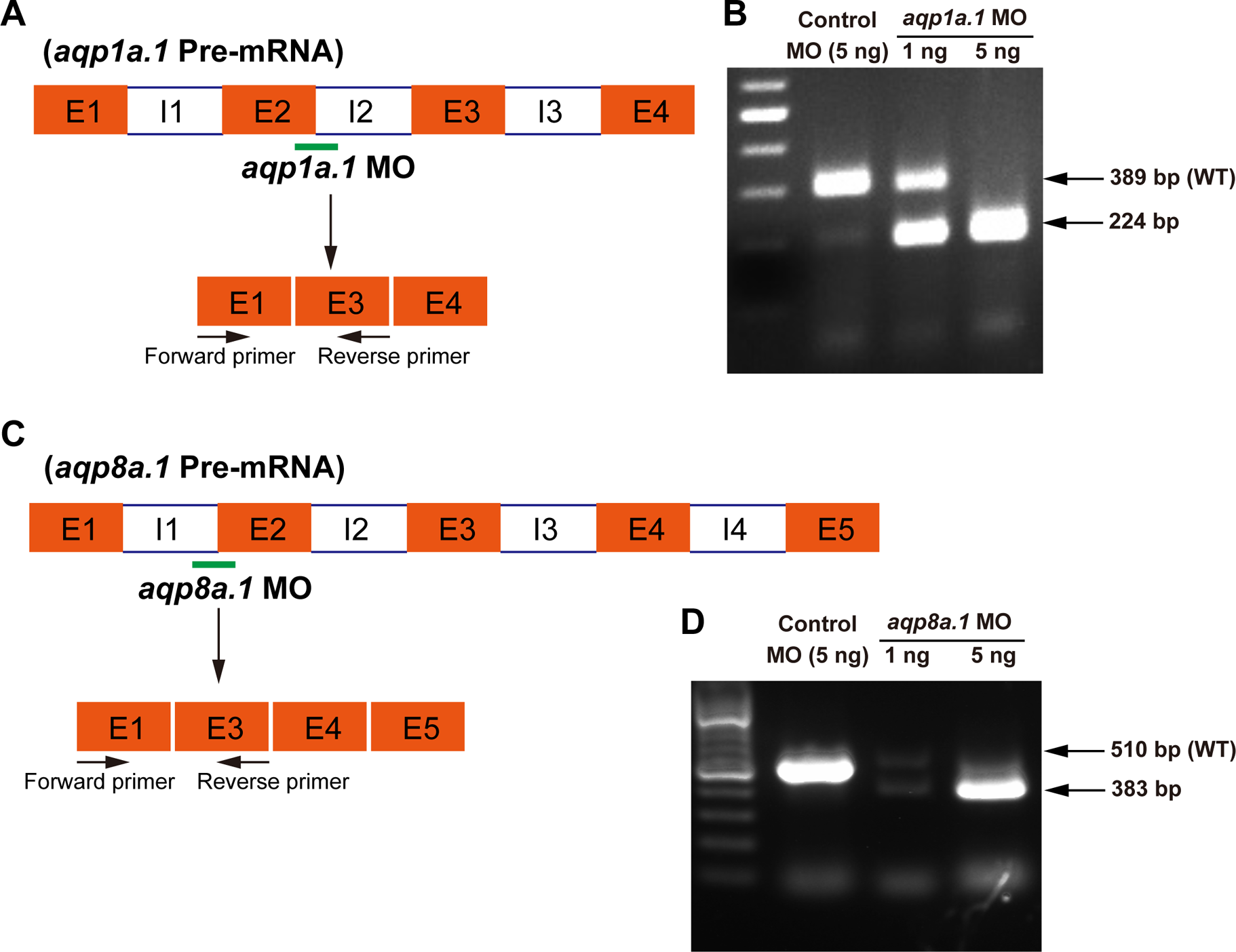
Validation of morpholino knock-down efficiency. **A,** Schematic representation of *aqp1a.1* splice blocking morpholino that targets the splicing site between the 2^nd^ exon and the 2^nd^ intron. RT-PCR primers to detect the presence or absence of exon 2 are indicated. **B,** Injection of *aqp1a.1* morpholino causes a 165 bp deletion, as indicated by RT-PCR. An amount of 5 ng *aqp1a.1* morpholino injection is able to induce a complete loss of mature *aqp1a.1* mRNA. **C,** Schematic representation of *aqp8a.1* splice blocking morpholino that targets the splicing site between the 1^st^ intron and the 2^nd^ exon. RT-PCR primers to detect the presence or absence of exon 2 are indicated. **D,** Injection of *aqp8a.1* morpholino causes a 127 bp deletion, as indicated by RT-PCR. An amount of 5 ng *aqp8a.1* morpholino injection is able to induce a complete loss of mature *aqp8a.1* mRNA.

**Supplementary Figure 9.**
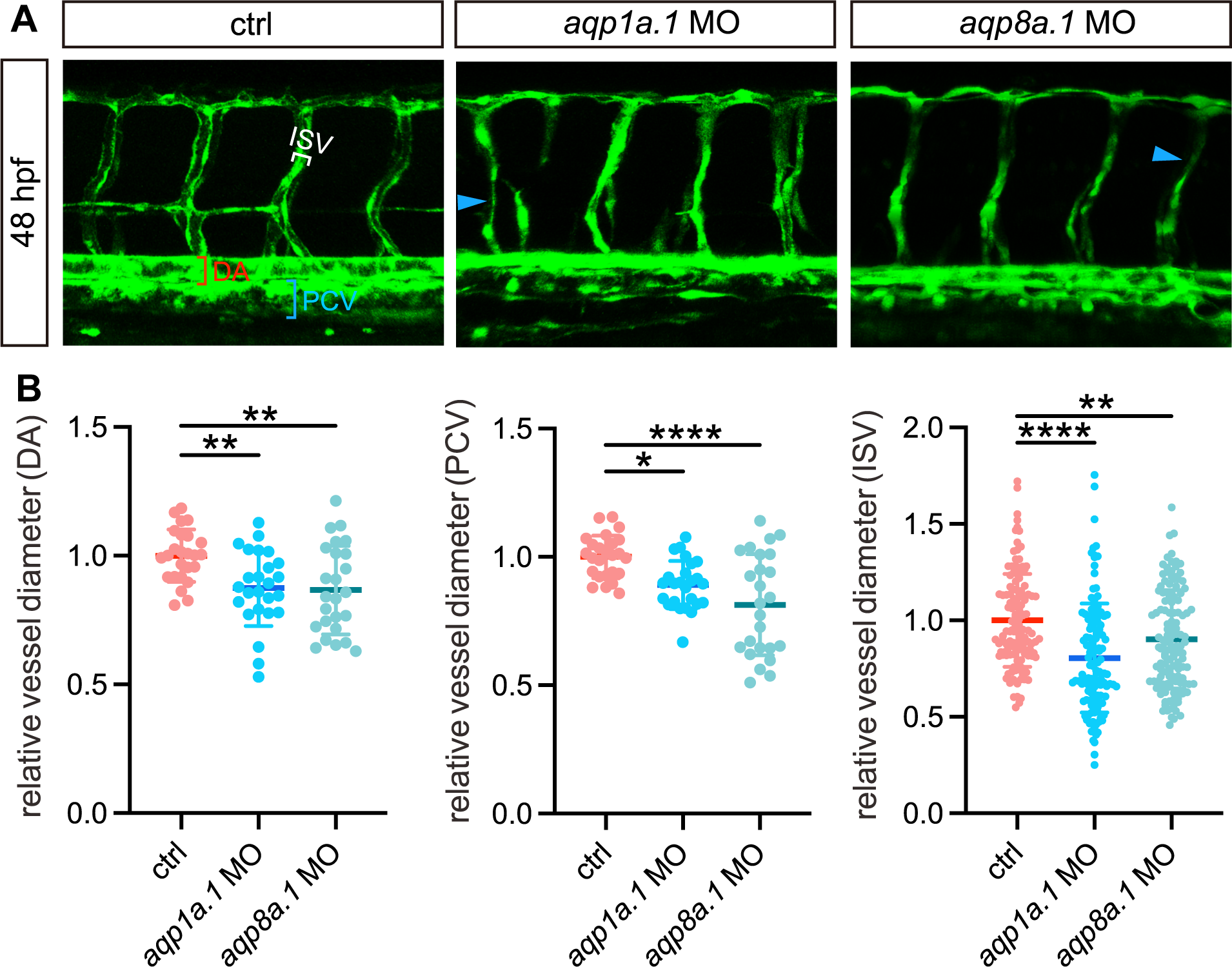
Knock-down of EC-enriched aquaporins (*aqp1a.1* or *aqp8a.1*) leads to the reduction of vascular diameter in zebrafish. **A,** Confocal images of ISV phenotypes in 48-hpf *Tg(kdrl:EGFP)* control embryos and *aqp1a.1*-knockdown and *aqp8a.1*-knock-down embryos. Red and blue square brackets indicate artery and vein, respectively. Blue arrowheads indicate the narrowed ISVs. **B,** Quantification of the diameters of ISV, DA, and PCV. Data are shown as mean ± s.e.m. One-way ANOVA analysis is applied. *, p<0.05. **, p < 0.01. ****, p < 0.0001.

**Supplementary Figure 10.**
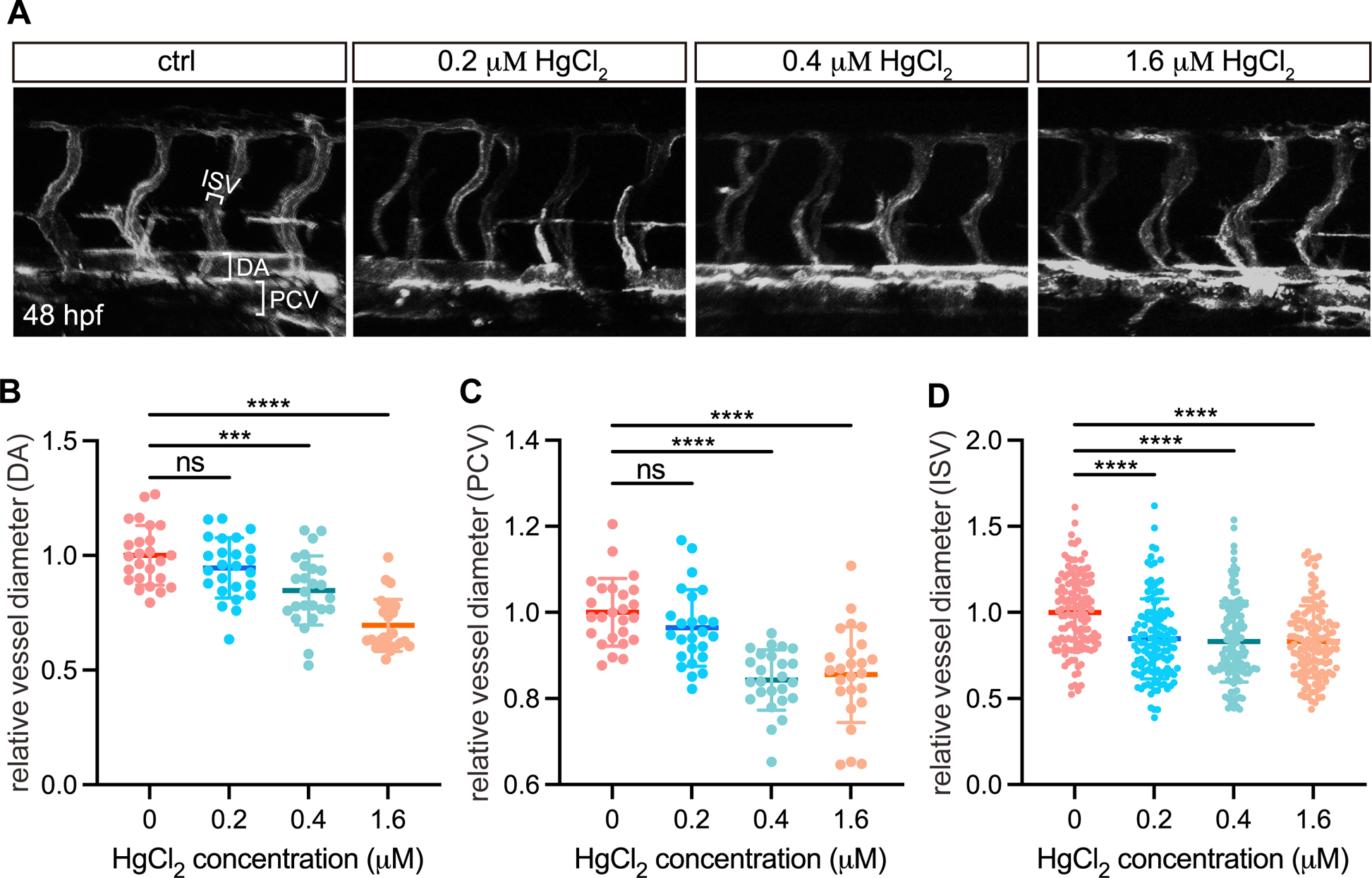
HgCl_2_ treatment narrows blood vessel caliber. **A,** Representative confocal images of ISV phenotypes in 48-hpf *Tg(kdrl:ras-mCherry)* upon HgCl2 treatment. **B-D,** Quantification of the diameters of ISV, DA, and PCV. Data are shown as mean ± s.e.m. One-way ANOVA analysis is applied. ns, not significant. ***, p < 0.001. ****, p < 0.0001.

**Supplementary Figure 11.**
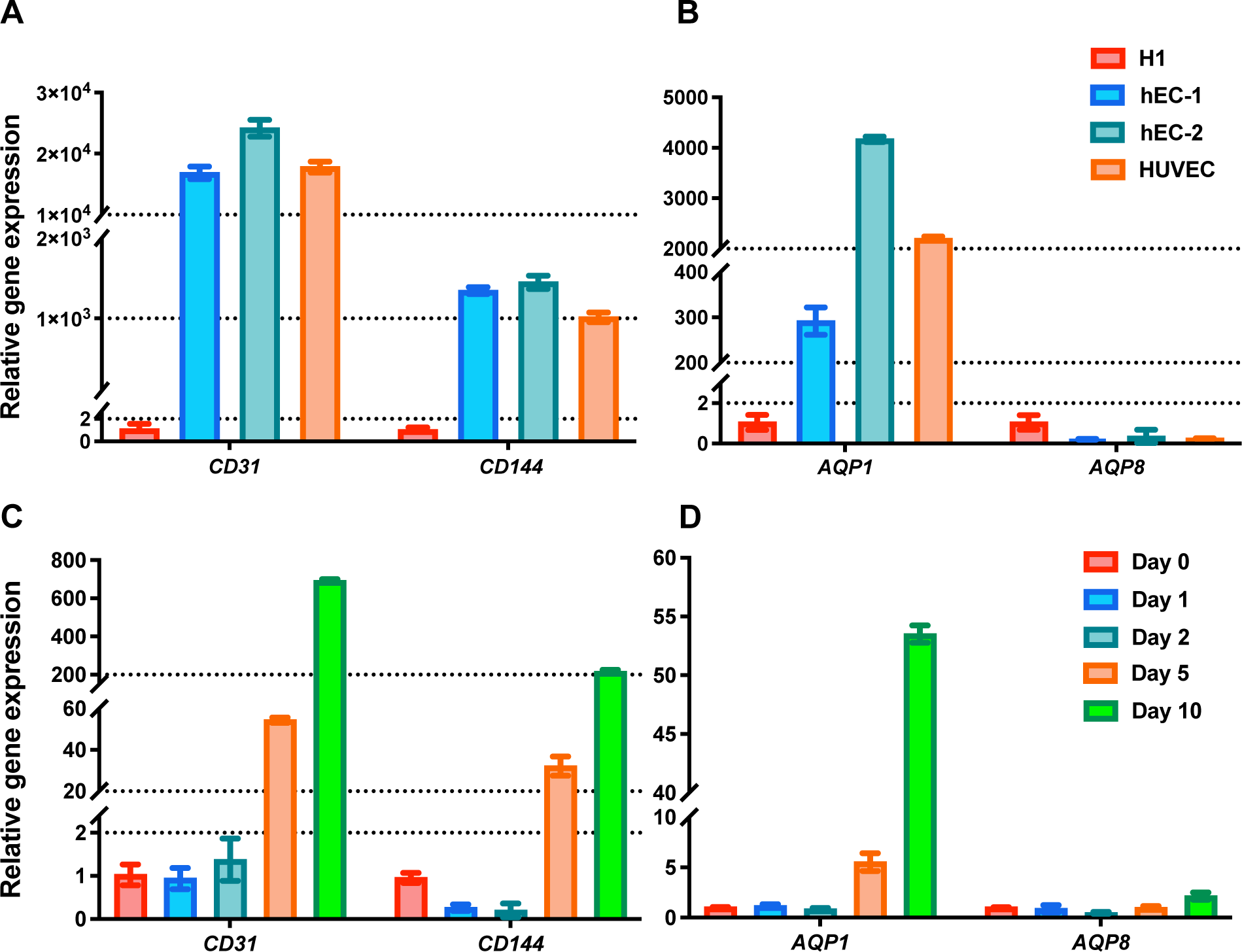
Q-PCR analysis of *CD31*, *CD144*, *AQP1,* and *AQP8* gene expressions. **A and B,** Relative gene expressions of *CD31*, *CD144*, *AQP1,* and *AQP8* in H1 stem cells, two human ECs (hEC-1 and hEC-2), and HUVECs, respectively. **C and D,** Relative gene expressions of *CD31*, *CD144*, *AQP1* and *AQP8* in H1 stem cell-derived ECs. EC differentiation is induced from day 4. Embryoid bodies for gene expression quantification are harvested on day 0, day 1, day 2, day 5, and day 10, respectively.

**Supplementary Table 1.**
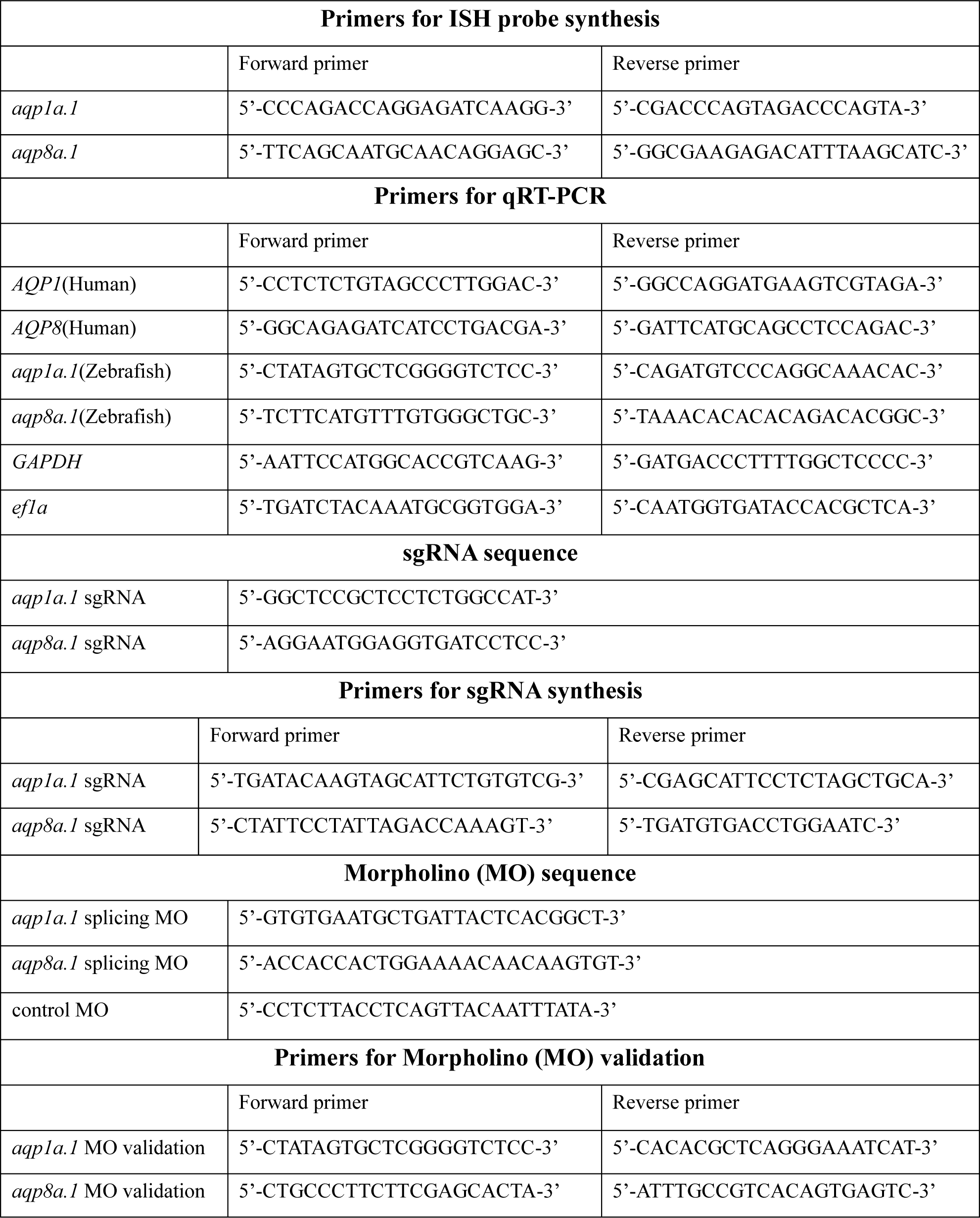
Primer list.

